# Quaternary ammonium compounds (QACs), QAC resistance genes, and QAC tolerant bacteria in livestock and human waste streams

**DOI:** 10.64898/2026.05.17.725718

**Authors:** Sophie Lennartz, Osas Elizabeth Aigbekaen, Alexandra Jahraus, Jan Siemens, Ines Mulder, Stefanie P. Glaeser

## Abstract

Quaternary ammonium compounds (QACs) are high production volume biocidal compounds increasingly scrutinized for their potential to promote antimicrobial resistance spread. This study compared the release of QACs, QAC resistance indicator genes (*qacE/qacEΔ1*), and QAC tolerant bacteria from livestock and human waste streams into the environment. Five livestock farms with on-farm biogas plants (BGPs), a rural and an urban municipal wastewater treatment plant (WWTP) were studied in parallel. In WWTPs, <1% of incoming QACs were discharged with treated wastewater but 10-20% were transferred to sewage sludge. QAC concentrations in sewage sludge far exceeded those in raw and digested manure. The *qacE/qacEΔ1* genes were detected in all samples with a higher relative abundance in solid than liquid samples. Relative abundances of QAC tolerant fast growing heterotrophic bacteria cultivated under high nutrient conditions at 37°C were higher in human than livestock waste streams. *Providencia* and *Pseudomonas* dominated the cultivated QAC tolerant bacteria in both systems but showed higher QAC tolerance when originating from human waste streams. Additionally, *Enterobacteriaceae* with higher QAC tolerance were cultivated from human waste streams. Most QAC tolerant strains carried antibiotic resistances without strong system differences. Only few strains carried the *qacE*/*qacEΔ1* gene indicating that other mechanisms must be responsible for the increased QAC tolerance. In conclusion, QACs, *qacE/qacEΔ1*, and viable QAC tolerant bacteria including potential pathogenic bacteria were released from livestock and human waste streams into the environment with highest abundances in a post-pandemic sewage sludge sample.

**Highlights:**

- QACs most abundant in human waste streams, especially biosolids
- Higher relative abundance of QAC tolerant bacteria in human waste streams
- *Pseudomonas* and *Providencia* dominated QAC tolerant bacteria in both waste streams
- *Enterobacteriaceae* with higher QAC tolerance abundant in human waste streams
- Most QAC tolerant strains carried additional antibiotic resistances

**Environmental implication:** Municipal wastewater treatment plants (WWTPs) and livestock farms are hotspots for antimicrobial resistance (AMR) propagation. We compared the simultaneous occurrence of quaternary ammonium compounds (QACs), resistance genes (RGs), QAC-tolerant bacteria, and their multidrug-resistance status in livestock and human waste streams. QACs, indicators of QAC tolerance and AMR occurred in both systems but were higher in WWTPs, especially sewage sludge. Our findings highlight the need for prudent disinfectant use and enhanced waste treatments to reduce the risks of spreading micropollutants, pathogens, and AMR via organic fertilizers or treated wastewater recycled in circular agricultural practice.

## 1. Introduction

Antimicrobial resistance (AMR) is one of the most urgent global public health threats of the 21st century [1–4]. The environmental dimension of AMR transmission and spread now receives increasing attention [5]. Beside pharmaceutical antibiotics, other biocidal compounds with antimicrobial properties can contribute to AMR via complex co- and cross-resistance mechanisms [6,7]. Compared to pharmaceuticals, disinfectants based on biocidal compounds are often applied in higher volumes and less target-specific [8,9], leading to potentially much higher emissions into the environment [10].

Quaternary ammonium compounds (QACs) are an important class of biocidal compounds and cationic surfactants, found among others in disinfectants, pesticides, detergents, and personal care products [11–14]. QACs carry a characteristic positively charged nitrogen atom surrounded by four covalently bound organic substituents, at least one of which is a hydrophobic alkyl chain of variable length. This amphiphilic structure determines the antimicrobial effects, surface-activity, and environmental behaviour of QACs [11,12,15,16]. Three common QAC subclasses can be differentiated: alkyltrimethyl-, benzylalkyl-, and dialkyldimethyl ammonium compounds (ATMACs, BACs, and DADMACs, respectively; for structural details see [12]).

QACs are effective against both, Gram-positive and Gram-negative bacteria, and thus frequently used in biocidal products [17,18]. Upon contact of QACs with microbial cells, electrostatic forces between the quaternary nitrogen and the negatively charged phospholipid bilayer of microbial cells can cause intercalation of the hydrophobic part (e.g. long alkyl chains) of the QAC molecules into the microbial membrane [17,19]. High QAC concentrations therefore induce membrane lysis followed by cell death. QAC concentrations below minimal inhibitory concentrations (sub-MIC values) can have different effects on bacterial cells including the loss of membrane osmoregulation and oxidative stress which triggers the SOS response and in consequence can increase DNA mutation rates and resistance gene transfer [20]. One of the most discussed resistance mechanisms of bacteria in the presence of high QAC concentrations is the induction of efflux systems including a broad range of efflux pumps determined either in Gram-positive or Gram-negative bacteria [21]. Often correlated with QAC tolerance in Gram-negative bacteria are small multidrug resistance (SMR) efflux pumps. Those non-specific efflux pumps can transport a wide range of toxic substances, including biocidal compounds such as QACs, but also antibiotics and heavy metals, out of bacterial cells [14]. The coding genes are often named as *qac* genes, e.g. *qacE*, *qacEΔ1* (functional *qacE* deletion variant with reduced activity [22], *qacG*, or *qacH* [23–25]). Especially *qacE/ qacEΔ1* is frequently used as a proxy to monitor and quantify the presence of QAC resistant bacteria in the environment [26–29]. It is controversially discussed if the presence of those genes is linked to an increased QAC tolerance [21,30]. But there is clear evidence that the presence of *qac* genes can be linked to the multi drug resistance (MDR) status of bacteria because *qac* genes, especially *qacE/ qacEΔ1* are often found on mobile genetic elements with other resistance gene (RGs) [26,31,32]. Those findings indicate that the combined dissemination of QACs and QAC tolerant bacteria into the environment can affect the transmission and proliferation of QAC tolerant and MDR bacteria via co- and cross-resistance mechanisms [12,33–36]. So far only a couple of studies selectively cultivated QAC tolerant bacteria from environmental samples using culture media supplemented with QACs [31,35]. QAC tolerant bacteria from livestock and human wastewater streams were not systematically studied yet, although these systems are among the most problematic sources of AMR transmission into the environment [37].

Up to roughly 75% of the annually used QACs enter wastewater treatment plants (WWTPs) [38–40], from where they are released into the aquatic and terrestrial environment via treated wastewater (WWTP effluent) discharged into rivers or biosolids applied as fertilizers onto agricultural fields [12,26,41]. QACs have already been studied in WWTPs [9,10,38,40,42–47] and wastewater-impacted matrices such as surface waters [48–50], suspended particulate matter [49,51,52], sediments [48,53–55], and long-term wastewater-irrigated soils [56]. Comparatively little is known about the emissions from animal husbandry. QACs can enter livestock waste after stable and equipment disinfection [41,57,58] and currently, about 30% of the applied veterinary disinfectants in Germany contain QACs, especially BACs and DADMACs [59]. To our knowledge, only two studies from the Netherlands and Japan have investigated QACs in livestock manure and wastewater [58,60] while two studies from Austria and Denmark reported the presence of QACs in BGP digestates [58,60]. A comparative investigation of the emissions of QACs and QAC tolerant bacteria from livestock and human waste streams is still missing.

The potential role of QACs in the environmental dimensions on AMR spread increased since the emergence of the severe acute respiratory syndrome coronavirus 2 (SARS-CoV-2) in March 2020. With the onset of the SARS-CoV-2 pandemic, disinfection practices increased drastically [61] and due to their antiviral properties, QACs were used as active component in a large fraction of the applied disinfectants [62,63]. In the USA, disinfectant use rose by 146% within the first year of the pandemic while German sales of QAC-based disinfectants increased by about 80% [64,65]. The annual production five years after the start of the pandemic is still exceeding the production in pre-pandemic years [66]. Alygizakis et al. [10] pointed out that QACs, including ATMACs, BACs, and DADMACs, showed the highest increase among different chemicals in incoming wastewater in Greece after the lockdown started (+331%). Tentative increases in the concentrations of these QACs have also been observed in suspended particulate matter of German rivers during the pandemic [51]. Thus, QAC concentrations, and consequently QAC-tolerance and multidrug resistance in WWTPs may have increased since the pandemic [64,67].

It is still not well understood under which environmental conditions released QACs are biologically active and can affect bacteria. QACs sorb strongly to organic matter and negatively charged clay mineral surfaces present in manure, sewage sludge, and soil [19,39,68–73]. This can affect their bioavailability to microorganisms and thereby both antimicrobial activity and biodegradability [74]. For example, Heyde et al. [68] have shown that sorption to and intercalation of QACs into 2:1 layer silicates buffers their acute toxicity to potentially pathogenic Gram-positive and Gram-negative bacteria found in livestock and human wastewater samples, while the presence of non-swellable 1:1 layer silicate did not. Especially under oxygen-limited conditions, sorption can also cause accumulation and long-term persistence of QACs as documented for processed sludge, soils, and sediments [11,16,41,48,56,75,76]. On the other hand, Mongelli et al. [45] found that QACs sorbed to the solid phase of activated sludge are still microbially degraded and thus accessible to microorganism. It is unclear if QACs accumulated in and on solid environmental matrices contribute to increasing QAC tolerance of bacteria attached to the same particles and consequently to horizontal gene transfer (HGT) and MDR formation.

This study investigated the combined abundance of QACs, QAC resistance indicator genes, and QAC tolerant bacteria in livestock and human waste streams including key output sources (manure, BGP digestates, biosolid, and treated wastewater) for pre and post pandemic sampling time points. The following hypotheses were investigated: (1) QACs, QAC RGs, and QAC tolerant bacteria are released from both, livestock and human waste streams into the environment. We expected a higher abundance of QAC tolerant bacteria in systems with higher QAC abundance (human waste streams). (2) Due to the strong sorption of QACs to organic material, higher concentrations of QACs are expected in solid compared to liquid wastewater and activated sludge fractions. If the particle-bound QACs remain biologically active, a higher abundance of QAC tolerant bacteria is consequently also expected in those waste stream samples. (3) Based on the current knowledge on co- and cross resistance mechanisms between QACs and antibiotics, we expected that QAC tolerant bacteria are also resistant to several classes of antibiotics.

To investigate these hypotheses, we analysed the occurrence of QACs, the QAC resistance indicator genes *qacE/qacEΔ1*, and quantified and characterized the fraction of culturable QAC tolerant bacteria from the same set of samples. Liquid and solid samples were collected from five livestock farms with small-scale on-farm BGPs and two municipal WWTPs in Germany. The studied livestock systems included dairy cow and pig farms with mesophilic or thermophilic BGPs processes. The human waste streams systems comprised one rural WWTP and one urban WWTP, which received the effluents of hospitals and a veterinary clinic. Two of the farms and the rural WWTP were studied both before and after the onset of the SARS-CoV-2 pandemic.

## 2. Material and methods

### 2.1 Livestock waste stream samples

Manure and anaerobic digestates were collected at five German farms with small-scale on-farm BGPs (see Table S1). The examined BGPs represented continuous stirred tank reactors with wet fermentation processes (dry mass <15% w/v). The liquid manure samples were taken from storage tanks, which collected manure as input material for the on-farm BGPs. Anaerobic digestates were taken from the post-fermenter storage tanks. At every sampling point, manure/slurry samples from the storage tank (BGP input material; BGP-I) and anaerobic digestates (BGP output samples; BGP-O) were taken simultaneously. Sampling campaigns were carried out between September 2019 and April 2020. Samples were taken at four time points from two of the farms (BGP 1 and 2) including pre- and post-SARS-CoV-2 pandemic, and at one time point from three farm (BGP 4, 5, and 6). Manure sources, BGP process parameters and sampling data are described in Table S1.

### 2.2 Human waste stream samples

Human wastewater samples were collected in two WWTPs, one rural (WWTP1) and one urban (WWTP6) (Table S2). Both WWTPs were in the same region of Germany. The rural WWTP used one activated sludge (AS) tank that shifted between aerobic and anaerobic phases. The collected primary and surplus sludge were combined and dewatered by centrifugation and not processed further. The urban WWTP operated with separate aerobic and anaerobic AS tanks and the collected sludge was anaerobically digested and dewatered. The following samples were collected for analyses: untreated wastewater (influent after mechanical treatment, WWTP-I), activated sludge (AS from the aerobic phase/tank), sludge (S, dewatered sludge, or ADS, anaerobically digested and dewatered sludge) and treated wastewater (final effluent released into the adjacent rivers, WWTP-E). The rural WWTP was sampled once before the onset of SARS-CoV-2 pandemic in November 2019 and twice after in May 2020 and March 2021. The urban WWTP was only sampled once in November 2021 (post-pandemic).

### 2.3 Sample collection and storage

All samples were placed in closed storage boxes containing ice packs immediately after sampling to ensure continuous cooling during the transport to the laboratory (approximately 4°C). Autoclaved 2 L glass bottles (SCHOTT, Germany) were used to collect samples for microbiological analyses. At the laboratory, samples were stored at 6°C and processed within 24 h after sampling. Samples for chemical analyses were collected in 500 mL and 250 mL polyethylene (PE) bottles (Corning, USA) and stored at -20°C until further analyses. Before freezing, the activated sludge samples were separated into solid and liquid phases by centrifugation (4,160 rpm, Rotana 460R, Hettich, Tuttlingen, Germany). Dry weights of all samples were determined by lyophilization (Table S3). Samples for molecular biological analysis were handled differently based on the samples type. Solid samples were collected in sterile 50 mL PE screw cap tubes (Greiner) and frozen immediately at -20°C; liquid samples were filtered to concentrate bacterial cells from larger water volumes using Sterivex-GP polyethersulfone filter units with a pore size of 0.22 µm (Merck Millipore, Darmstadt, Germany). A serial filtration was used for some samples to separate particle-attached and free-living bacteria. Particle-attached bacteria were either collected on nitrocellulose filter membranes (8 µm pore size, 47 mm diameter; Sartorius, Germany) or on sterile Minisart NML surfactant-free cellulose acetate (SFCA) syringe filters (5 µm pore size; Sartorius). The remaining free-living bacteria were collected from the flow-through of larger particle size filters on Sterivex-GP 0.22 µm filter units. The remaining water was removed from filter units before they were stored at -20°C until DNA extraction.

### 2.4 Extraction and analysis of QACs

#### 2.4.1 Standards and chemicals

QAC reference standards were purchased from TCI, Sigma Aldrich and HPC Standards and had a chemical purity of at least 95% (Table S4). For routine analysis, a multi-standard of all target analytes and two internal standard (IS) mixtures of deuterated QACs were prepared in acetonitrile. The QAC multi-standard contained even-chained ATMACs of length C8 to C16, BACs C8 to C18, DADMACs C8 to C18, benzethonium chloride (BEC) and chlormequat chloride (CLQ). IS mixture A comprised D_7_-labelled BACs C8, C12, C14, C18, and D_6_-labeled DADMACs C8 to C14. IS mix B was similar but did not include BAC-C14-D_7_ and DADMAC-C14-D_6_. HPLC grade acetonitrile, methanol, ethyl acetate, and formic acid for QAC extractions were purchased from VWR (Darmstadt, Germany). Pentane was obtained from Carl Roth (Karlsruhe, Germany). Ammonium formate was bought from Acros Organics, and 30% (w/w) ammonium hydroxide solution and 35% (w/w) hydrochloric acid from Merck (Darmstadt, Germany). Ultrapure “milli-Q” water was prepared in a purelab flex system (Elga LabWater, Germany).

#### 2.4.2 QAC extractions

Depending on the sample matrix, different QAC extraction procedures were employed. For manure, BGP digestates, and sewage sludges, an ultrasonic-assisted extraction with subsequent solid-phase extraction (SPE) with chromabond CN cartridges (Macherey-Nagel, Düren, Germany) originally developed by Heyde et al. [77] was applied with minor modifications using 1.5-2 g of freeze-dried sample material. The same ultrasonic extraction but without the SPE was used for the particulate phase of WWTP influent and effluent separated from the liquid phase by centrifugation (3,850 g for 30 min). The liquid phase of the WWTP influent, activated sludge and effluent was processed by SPE over Oasis HLB 150 cc cartridges (Waters) using 150-200 mL sample aliquots based on a protocol adapted from Östman et al. [9]. Total QAC concentrations were obtained by summing concentrations in both liquid and solid fractions expressed in ng L^-1^ based on the bulk sample volumes. The particle-associated concentrations in activated sludge were initially determined in ng/g dry weight and then converted to liquid-phase concentrations based on the dry matter content given in the Supplementary Data. All samples were extracted in triplicate when sufficient material was available, however, this was not always possible due to the retrospective analysis. Detailed description of the extraction procedures is given in Text S1 in the supplementary information. The performance of each extraction method was assessed in terms of recovery (Tables S5-S6), matrix effect (Tables S7-S9), and method detection and quantification limits (MDL and MQL) as described in more detail in Text S2.1-S2.3 in the supplementary information. Notably, SPE resulted in very low recoveries for DADMAC-C18 and CLQ. These homologues were therefore not included in the calculation of ∑DADMAC and ∑QAC concentrations.

#### 2.4.3 QAC quantification by high performance liquid chromatography-tandem mass spectrometry (HPLC-MS/MS)

All extracts were fortified with one of the IS mixtures to 20 ng mL^-1^ (final concentration) of each IS prior to measurement to correct for instrumental drift and matrix effects. QACs were analysed with a Sciex Exion LC™ system coupled to a Sciex Qtrap 4500 tandem mass spectrometer operating with positive mode electrospray ionization. An XSelect CSH Phenyl-Hexyl-Column (130 Å, 150 mm length, 2.1 mm ID, 3.5 μm particle size, Waters) with corresponding guard column was used for chromatographic separation. The mobile phases consisted of a buffer solution with 50 mM formic acid and 10 mM ammonium formate (A), and acetonitrile (B). Extract aliquots of 20 µL were injected at a mobile phase flow rate of 0.25 mL min^-1^ and QACs were eluted with the following gradient profile: 0 – 3 min: increase from 0% to 75% B, 3 – 14.5 min: increase to 100% B, 14.5 – 21 min: isocratic hold at 100% B, 21 – 26 min: switch to 100% A. The analytes were measured in multiple reaction monitoring (MRM) mode and quantified by external linear calibration (no weighting) using the analyte to IS area ratio over a concentration range of 0.5 – 200 ng mL^-1^. The linear range was assessed individually for each QAC homologue and extracts were diluted between 10 to 1000 times with acetonitrile if necessary. Presence of the qualifier ion was required for positive identification. As a default, the ion ratio tolerance was set to 30%, followed by visual confirmation on case-by-case basis using Sciex OS v. 2.0.0. Further details on the MRM transitions, source parameters and method performance are provided in the supplementary information (Tables S10-S12; Text S2.3-S2.5).

### 2.5 Extraction of total DNA from waste stream samples

Total DNA of solid and liquid wastewater samples was extracted with the NucleoSpin Soil DNA extraction kit (Macherey Nagel). Either 0.3 to 0.5 g solid sample material or filter matrices containing bacterial cells collected from defined volumes of liquid sample material were used for DNA extraction. Filter units were pre-cooled in liquid nitrogen and opened with a hammer. A sterile scapula and forceps were used to transfer the bacterial cells containing filter material to the extraction tubes. Extraction was performed based on the protocol provided by the manufacturer with few modifications as described by Glaeser at al. [78]. The lysing buffer SL1 and 150 µL Enhancer SX solution were used during mechanical treatment in a Retsch mill MM400 (Retsch, Germany) with a frequency of 30 beats s^-1^ for 5 min. The final DNA elusion step was performed in two steps twice with 75 µL PCR grade water (Roth, Germany) preheated to 80°C. The two flowthroughs containing DNA were combined subsequently and stored at - 20°C. The DNA extracts were 1:30 diluted for subsequent PCR based analysis.

### 2.6 Quantification of the bacterial 16S rRNA gene, a mobile genetic element gene, and resistance genes (RGs)

Quantitative PCR was used to determine the abundance of bacterial 16S rRNA gene copies (to estimate bacterial abundance), *intI1* (integrase gene, marker gene for mobile genetic elements and anthropogenic pollution, [79]), and some RGs including *qacE*/*qacEΔ1* (QAC resistance indicator gene), and *sul1*, *sul2*, *tetM,* and *qnrS* (sulfamethoxazole, tetracycline, and fluoroquinolone RGs). Analysis was done according to previous studies [80,81] in a 10 µL volumes using a CFX100 Cycler (BioRad). The SsoFAST EvaGreen Supermix (Bio-Rad Laboratories, Feldkirchen, Germany) and the Applied Biosystems TaqMan Gene Expression Supermix (ThermoFischer Scientific, USA) were used for EvaGreen and TaqMan based qPCRs, respectively. Primer systems, standards, and qPCR conditions are given in Tables S13 and S14.

### 2.7 Cultivation and characterization of QAC tolerant bacteria

Bacteria were detached from 10 g fresh manure, BGP digestate, and dewatered sludge using 90 mL filter-sterilized 0.2% (w/v) tetra-sodium-pyrophosphate (TSPP) (Merck, Germany) buffer. Samples were added to TSPP in a sterile stomacher plastic bag (Seward Model 80, Germany) and treated in a Stomacher (Seward, Germany) for 2 minutes at maximum speed (300 rpm). A fraction of 40 mL of the bacterial suspensions was transferred to autoclaved glass bottles (SCHOTT, Germany). The cell suspensions were serially diluted up to 10^-7^ in autoclaved 0.9% (w/v) sodium chloride (NaCl, Merck, Germany) solution (total volumes 5 mL). Liquid samples, influent, activated sludge, and effluent samples, were directly serially diluted in autoclaved 0.9% (w/v) NaCl solution (total volumes 5 mL). From each dilution, 100 μL were plated per agar plate using the spread plate technique.

Fast growing heterotrophic bacteria and the fraction of fast growing heterotrophic QAC tolerant bacteria were cultivated on Mueller Hinton (MH) agar (Sigma-Aldrich, Germany) and MH agar supplemented with BAC-C12 (TCI, Japan) at 37°C. BAC-C12 was added in two concentrations, 50 and 100 µg BAC-C12 mL^-1^ (MH+BAC50 and MH+BAC100), after autoclaving to pre-cooled agar media. For this, a stock solution of 40 mg BAC-C12 in 8 mL of autoclaved pure water (5,000 µg BAC-C12 mL^-1^) was prepared and sterilized by filtration using a 0.45 µm syringe filter. The stock solution was short term stored in ice or frozen at -20°C until further use. Aliquots of this stock solution (2 and 4 mL) were added to 1 L pre-cooled autoclaved agar media, to obtain a final concentration of 50 and 100 µg BAC-C12 mL^-1^, respectively, in subsequently poured agar media plates. All agar plates were incubated under oxic conditions in darkness at 37°C for 24 to 48 h. Concentrations of colony forming units (CFUs) were calculated per g of solid and per mL of liquid samples. Standard deviations were set at 10% of the CFU counts.

### 2.8 Selections of abundant cultivated bacterial strains for detailed characterization

For detailed characterization of abundant cultivated bacteria, 2 to 8 colonies of different morphologies, were selected from the highest positive dilutions per medium and sample. Bacterial strains were purified following the 13-streak method using several transfer steps of single colonies until a pure culture was obtained. For long-term preservation of the living biological material two loops of fresh biomass of each pure culture were resuspended in 500 µL Gibco new-born calf serum (NBCS, Thermo Fisher Scientific) and stored in 1.4 mL U-bottom push cap tubes (Micronic, Netherlands) at -20 or -80°C, respectively. In parallel, a cell lysate was generated for each culture for subsequent molecular characterization. One inoculation loop of fresh biomass was suspended in 500 μL pure water (DNase and RNase free, PCR grade, ROTH, Germany) and treated with three cycles of freezing (−20°C) and thawing (1 min 50 seconds, 100°C, heating block) according to Schauss et al. [82]. Cell lysates were stored at -20°C.

### 2.9 Strain-level differentiation of cultivated bacteria by genomic fingerprinting

All strains were analysed using BOX-PCR with primer BOX1A (5’-CTACGGCAAGGCGACGCTGACG-3’) [83]. For strains without good BOX-PCR results, (GTG)_5_-PCR with primer GTG5 (5’-GTGGTGGTGGTGGTG-3’) was performed in addition [84] All analyses were performed according to Bartz et al. [85]. Comparative analysis of the BOX and GTG5 fingerprint patterns obtained after 1.4% (w/v) agarose electrophoresis and ethidium bromide staining was performed in Bionumerics 8 (bioMérieux, France). The unweighted pair group method with arithmetic mean (UPGMA) was used for clustering based on a distance matrix calculated with the Pearson’s correlation coefficient considering the presence/absence and intensity of the individual DNA bands. Each individual fingerprint pattern was assigned as a new genotype. Genomic fingerprint data were used to avoid repetitive testing of the same environmental bacterium (clonal strains) from individual samples.

### 2.10 Phylogenetic identification of cultivated bacteria by partial 16S rRNA gene sequencing

Representative strains of each genotype were phylogenetically identified by partial 16S rRNA gene sequencing. The nearly full-length 16S rRNA gene was amplified with the universal 16S rRNA gene targeting primers EUB9F (5’-AGAGTTTGATCMTGGCTCAG-3’) and EUB1492R (5’-GGYTACCTTGTTACGACTT-3’) [86] and sequenced by the Sanger sequencing technology by LGC Genomics (Berlin, Germany) using primer EUB9F as described in detail in Schauss et al. [82]. Sequences were corrected manually based on electropherograms in MEGA11 [87]. Unclear 5’ and 3’ ends were removed from Sanger sequences and the internal sequence quality was checked using the electropherograms and by the comparison of the sequences in an alignment with the next 10 related strains using the alignment option in ARB-Silva which aligned the sequence against the ARB seed alignment (https://www.arb-silva.de/aligner/). The ACT window was used to search and classify the sequences, with a minimum identity with query sequence of 0.95 and number of neighbours per query sequence limited to 10. For phylogenetic classification, the Basic Local Alignment Search Tool (BLAST) version blastn blast+ v2.13.0 of the NCBI (https://blast.ncbi.nlm.nih.gov/) was used to determine sequence similarities to the next related type strains by using the curated 16S ribosomal RNA type strain database (updated as of 2023/01/12). The genus-level assignment was performed if strains shared a partial 16S rRNA gene sequence similarity of at least >99.0% to type strain sequences representing species of the respective genus. This was the case for all studied strains.

All 16S rRNA gene sequences were submitted to GenBank and stored under Accession numbers OQ160434 to OQ160750.

### 2.11 Determination of minimal inhibitory concentrations (MICs) for BAC-C12 and DADMAC-C10

The minimal inhibitory concentrations (MICs) for BAC-C12 and DADMAC-C10 were determined for all strains excluding clonal strains obtained from one agar plate. Clinical Laboratory Standards Institute (CLSI) guidelines were followed for strain preparation and treatment to test for susceptibility against BAC-C12 and DADMAC-C10 [88]. Initial stock solutions of BAC-C12 and DADMAC-C10 were prepared with a concentration of 200 µg mL^-1^. The DADMAC-C10 stock solution was further ten-fold diluted in pure water. Stock solutions were filter-sterilized using 0.45 µm syringe filters. Serial dilutions for each compound were prepared using autoclaved pure water. MIC testing was performed in 96 well plates (Greiner Bio-One, Germany) as described by Heyde et al. [68]. Each well was pre-filled with 25 µL MH broth and 25 µL four-fold-concentrated QACs. Strains were cultivated overnight on Columbia agar with 5% sheep blood (Oxoid, ThermoFischer Scientific, USA) at 37°C. Fresh biomass was suspended in 0.9% NaCl to a McFarland Standard density of 0.5 and used for the inoculation of MH broth. An aliquot of 100 µL of this suspension was used to inoculate individual wells of the 96 well plates (100 µL per well). The tested concentration included 200, 100, 50, 25, 12.5, 6.25, and 3.125 µg BAC-C12 mL^-1^ and 20, 10, 5, 2.5, 1.25, 0.625, and 0.3125 µg DADMAC-C10 mL^-1^. The plates were closed with sterile sealing tape (Thermo Scientific, USA) and incubated for 24 h at 37°C in a dark, humid chamber. Growth was determined by visual examination. The lowest concentration that inhibited growth of a strain represented the MIC value.

### 2.12 Screening for *qacE*/*qacEΔ1* and *intI1* genes in QAC tolerant cultivated strains

Cell lysates of cultivated strains were used to screen for the presence of *qacE*/*qacEΔ1* and *intI1* genes. The above-described qPCR systems were used for these screenings.

### 2.13 Antimicrobial susceptibility testing (AST) and multiple antibiotic resistance (MAR) index determination for cultivated QAC tolerant strains

Bacterial strains cultivated on MH in the presence of BAC-C12 underwent antimicrobial susceptibility testing (AST) against 8 antibiotics using the disc diffusion methodology. Analysis was performed based on the EUCAST guidelines [89]. The following antibiotics were tested (concentrations in µg are given in brackets): piperacillin (PIP, 100), cefotaxime (CTX, 30 or 5), ceftazidime (CAZ, 10), ciprofloxacin (CIP, 5), meropenem (MER, 10), fosfomycin (FOS, 200), amikacin (AMK, 30), chloramphenicol (CMP, 30), tetracycline (TET, 30), gentamicin (GEN, 10), trimethoprim/sulfamethoxazole (TMP/SMX, 1.25/23.75). To test the Extended Spectrum beta-lactamase (ESBL) phenotype CTX (30) with clavulanic acid (CLA) was used in addition to CTX (30). A confirmed ESBL phenotype was obtained if the resistance to CTX was retained in the presence of CLA. Zone diameters were determined based on EUCAST (v14.0, 2024) and CLSI (M100-S30, [88]) breakpoint tables. Strains were classified as sensitive (S) or resistance (R) for each tested antibiotic. The multiple antibiotic resistance (MAR) index for individual bacterial strains was calculated as the ratio between the number of antibiotics the strain was resistant to, and the total number of antibiotics tested [90]. Mean MAR indices were further determined at the genus level by the calculation of a mean MAR index based in the MAR indices of individual strains assigned to the genus. Mean MAR indices were also separately calculated for groups of strains of a genus derived from different isolation sources.

### 2.14 Data analyses and statistics

QAC concentrations were determined by replicate extractions when sufficient material was available. Due to the retrospective analysis the number of extracted replicates differed between samples and is provided in the supplementary data file. Sample concentrations were calculated as the mean of all extracted replicates. Mean concentrations below the instrument quantification limit or MDL were treated as below detection limit (<LOD) and set to 0 for the calculation of sum parameters.

Nonparametric tests were used to investigate differences in QAC concentrations, the abundances of RGs, as well as total and QAC-tolerant culturable heterotrophic bacteria. Wilcoxon rank sum tests (2-sample comparison) were used for comparisons between the waste systems, input vs. output samples, and pre vs. post pandemic timepoints. Different process stages within WWTPs and the four different output samples from both systems (manure, BGP digestates, digested sludge/biosolids, effluent) were compared by Kruskal-Wallis test with Conover-Iman post hoc comparison. Spearman correlation coefficients were calculated between QAC concentrations, the abundances of *qacE*/*qacEΔ1, intl1*, and antibiotic RGs, and the proportion of QAC-tolerant bacteria. Variables with >50% of all concentration values <LOD were omitted from the correlation analysis. Obtained p-values were adjusted for multiple testing using the Benjamini-Hochberg procedure. In all tests statistical significance was evaluated at p < 0.05. R v.4.3.3 [91] and SigmaPlot version 15 (Applied Maths) were used for data analysis and graphical presentation.

## 3 Results

### 3.1 QAC concentrations in the studied waste stream systems

Samples from both, livestock and human waste streams, contained all three major QAC subclasses, ATMACs, BACs, and DADMACs (Fig. 1, Table S15). While extraction efficiencies and detection limits varied for different matrices and homologues, QACs were overall much more abundant in the human compared to livestock waste streams.

**Fig. 1.**
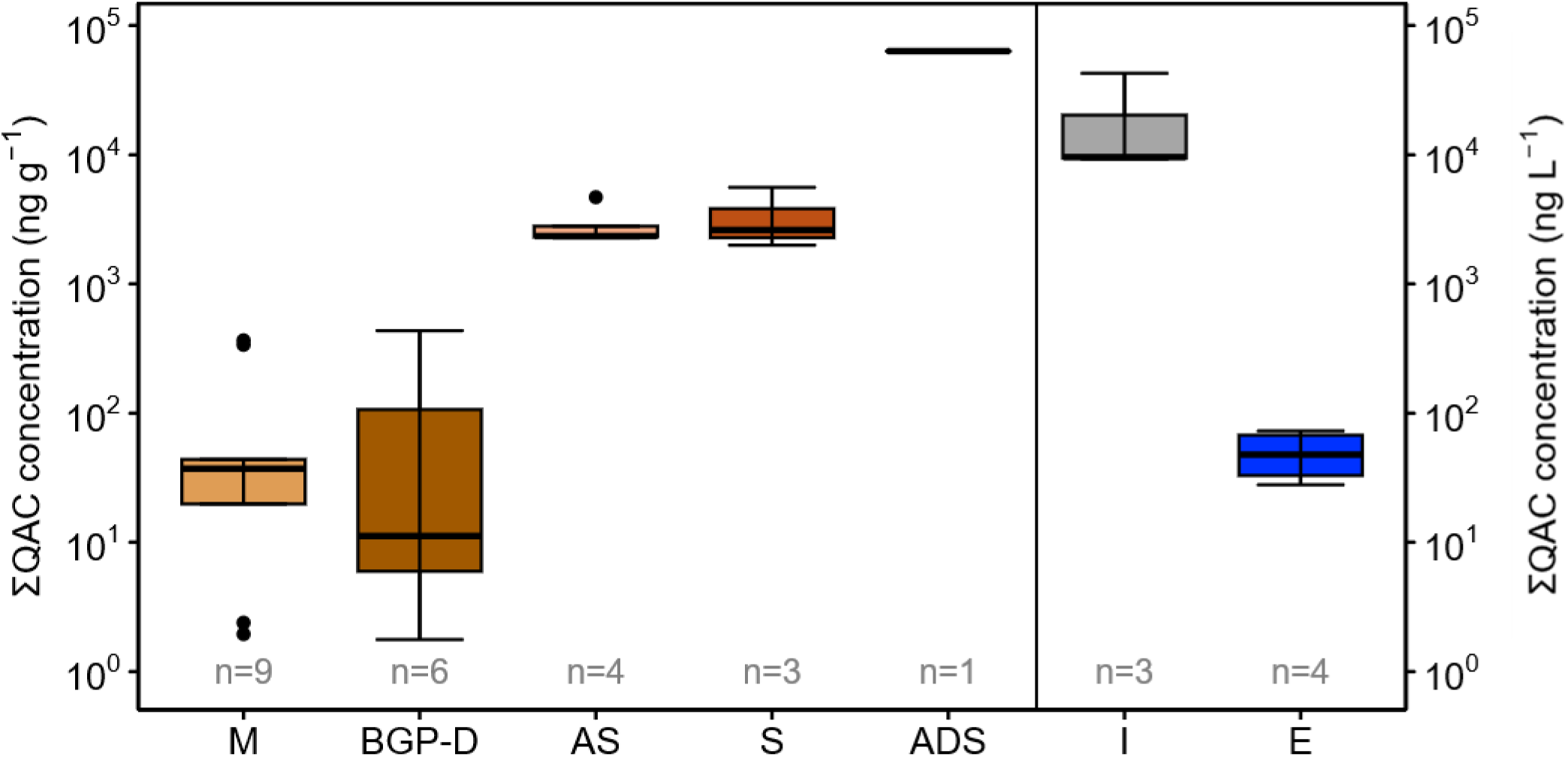
Total QAC concentrations > LOD in livestock and WWTP system. The number of samples with concentrations > LOD is shown by n (replicate analyses averaged into one value per sample). For WWTP samples, n is equal to the number of investigated samples. For the livestock systems, 11 manure and 11 BGP-D samples were investigated. M: manure; BGP-D: digested manure; AS: activated sludge; S: dewatered sludge; ADS: anaerobically digested sludge; I: influent; E: effluent.

#### 3.1.1 Livestock systems

QACs were detected in 10 out of 11 raw manure samples with ∑QAC concentrations ranging from <LOD to 365 ng g^-1^ (median 25 ng g^-1^). Concentrations were significantly higher (p_adjusted_ = 0.002, one-sided Wilcoxon rank-sum test; see Table S16) for pig manure (337-365 ng g^-1^) compared to cattle manure (<LOD-44 ng g^-1^). The highest concentrations were measured for BAC-C12 (285 ng g^-1^), DADMAC-C10 (247 ng g^-1^), and BAC-C14 (51 ng g^-1^). Only BACs C12-C14 were detected at all farms. The other homologues, ATMACs C12-C14, BAC-C10, BAC-C16, and DADMAC-C8, only occurred in 1-2 samples each. After anaerobic digestion, QACs were only detected in 6 out of 11 BGP digestate samples. The ∑QAC concentrations were overall slightly, though not significantly, lower than in the raw manure (<LOD to 433 ng g^-1^, median 2 ng g^-1^; see Table S17 for detailed statistical results). However, in BGP-2, ATMAC-C16 was only detected in the digestates while in BGP-4, higher concentrations of ATMAC-C10 and BACs C12-C14 were measured in digestates than raw manure.

#### 3.1.2 Human-wastewater treating WWTPs

∑QAC concentrations in the untreated human wastewater (WWTP influent) amounted to 9.3 - 9.6 µg L^-1^ in the rural WWTP and 42 µg L^-1^ in the urban WWTP. In contrast to manure, 14-16 out of 17 quantified QACs were found above the LOD. The most abundant homologues were BACs C12 -C14, DADMACs C8 -C10, and ATMAC-C16. DADMAC-C10 alone made up to 32-58% of the ∑QACs and BAC-C12 12-43% (Fig. S1). In both WWTPs, the concentrations of ∑DADMAC and ∑BACs exceeded ∑ATMACs. Between 71-81% of ∑QACs in incoming wastewater were measured in the particle-associated phase (see Fig. S2; Table S18). This distribution was similar for all QAC subclasses (∑ATMACs: 63-76%, ∑BACs: 71-82%, ∑DADMACs: 73-82%), but a notable increase in solid-phase partitioning was observed with increasing chain length, especially for ATMACs (Fig. S2; Table S18).

The effluent wastewater concentrations of ∑QAC ranged from 28 to 73 ng L^-1^ in both WWTPs, corresponding to strong reduction in ∑QAC concentrations of over 99% (76-100% for individual homologues) compared to the influent (p = 0.03; p_adjusted_ = 0.09; one-sided Wilcoxon rank sum test; Table S19). Only few QACs, including ATMACs C8-C10, BAC-C12, and DADMACs C8-C10 were detected in the effluent. While DADMACs were found in both, the particle-associated and liquid phase, BAC-C12 was only detected particle-associated and ATMACs only in the liquid phase (see Table S18).

∑QAC concentrations in activated sludge were in the same order of magnitude as in the influent, ranging between 12-17 µg L^-1^ in the rural WWTP and 27 µg L^-1^ in the urban WWTP (Table S18). In both WWTPs, activated sludge showed a similar homologue distribution as the influent, albeit with a slightly higher share of DADMACs and particularly DADMAC-C10 (43-71% of ∑QACs; Fig. S1). Particle-associated QACs comprised 70-100% of the total concentrations in activated sludge for all homologues except ATMAC C8, which was only found in the liquid phase (Fig. S2; Table S18). The corresponding solid phase concentrations ranged from 2.3 to 4.7 µg ∑QACs g^-1^ dw (Fig. 1).

∑QAC concentrations in the dewatered sludge of the rural WWTP fell in the same range (2.0 to 5.6 µg g^-1^) and showed a similar homologue distribution as the activated sludge (∑DADMACs > ∑BACs > ∑ATMACs). DADMAC-C10 alone comprised 66-71% of ∑QACs. In contrast, in the urban WWTP, ∑QAC concentrations in the anaerobically digested sludge were over 10-fold higher than in activated sludge, amounting to 63 µg g^-1^. BAC-C12 and DADMAC-C10 made up the largest fraction of 49% and 29% of ∑QAC (Fig. S1), respectively. Based on the annually treated wastewater volume and sludge production in both WWTPs for the year 2021 (see Table S2), an estimated 8-22% of the incoming QACs have thus not been degraded and ended up in the sewage sludge (Fig. S3).

#### 3.1.3 Comparative QAC emissions from both waste stream systems

Considering the above data and the nation-wide production of sewage sludge and wastewater in 2019-2020 [92,93], estimated annual QAC mass flows into sewage sludge (up to 110 t/a) far exceeded those with WWTP effluent (<1 t/a). When comparing the different types of potential biofertilizers from the human and livestock waste streams, digested sewage sludge contained significantly higher concentrations of BAC-C12, BAC-C14, DADMAC-C10, ∑ATMACs, ∑BACs, ∑DADMACs and ∑QACs than raw and digested manure (p < 0.05, Kruskal-Wallis tests with Conover-Iman post hoc comparison; Table S17). Consequently, estimated QAC inputs into German soils based on the annual application volumes of each organic fertilizer [92,94] are around 6-fold to 11-fold higher for digested sewage sludge (16 t/a) compared to BGP digestates (2.3 t/a) or raw liquid manure (1.4 t/a).

### 3.2 Bacterial abundance in studied wastewater streams

The abundance of *Bacteria* in studied livestock and human waste stream samples was estimated by the quantification of bacterial 16S rRNA gene copies in total extracted DNA. The concentrations were in the range of 10^10^ to 10^11^ 16S rRNA gene copies per g in livestock samples (manure, BGP digestates) as well as in activated sludge and sludge samples of the rural and urban WWTPs (Fig. 2A, Table S15). The concentrations were overall slightly lower in WWTP sludges, and significantly lower in activated sludge than livestock samples (p < 0.05; Kruskal-Wallis test with Conover-Iman post-hoc comparison; Table S17). The concentrations in untreated human wastewater (WWTP influent) were in the range of 10^7^ to 10^9^ gene copies per mL, while the concentrations in treated wastewater (WWTP effluents) were strongly reduced by 2 to 3 log scales (p = 0.03; p_adjusted_ = 0.09; one-sided Wilcoxon rank sum test; Table S19) compared to untreated wastewater. This corresponds to an elimination rate of 98.4 to 99.9% of the bacterial load of untreated wastewater by the wastewater treatment processes.

**Fig. 2.**
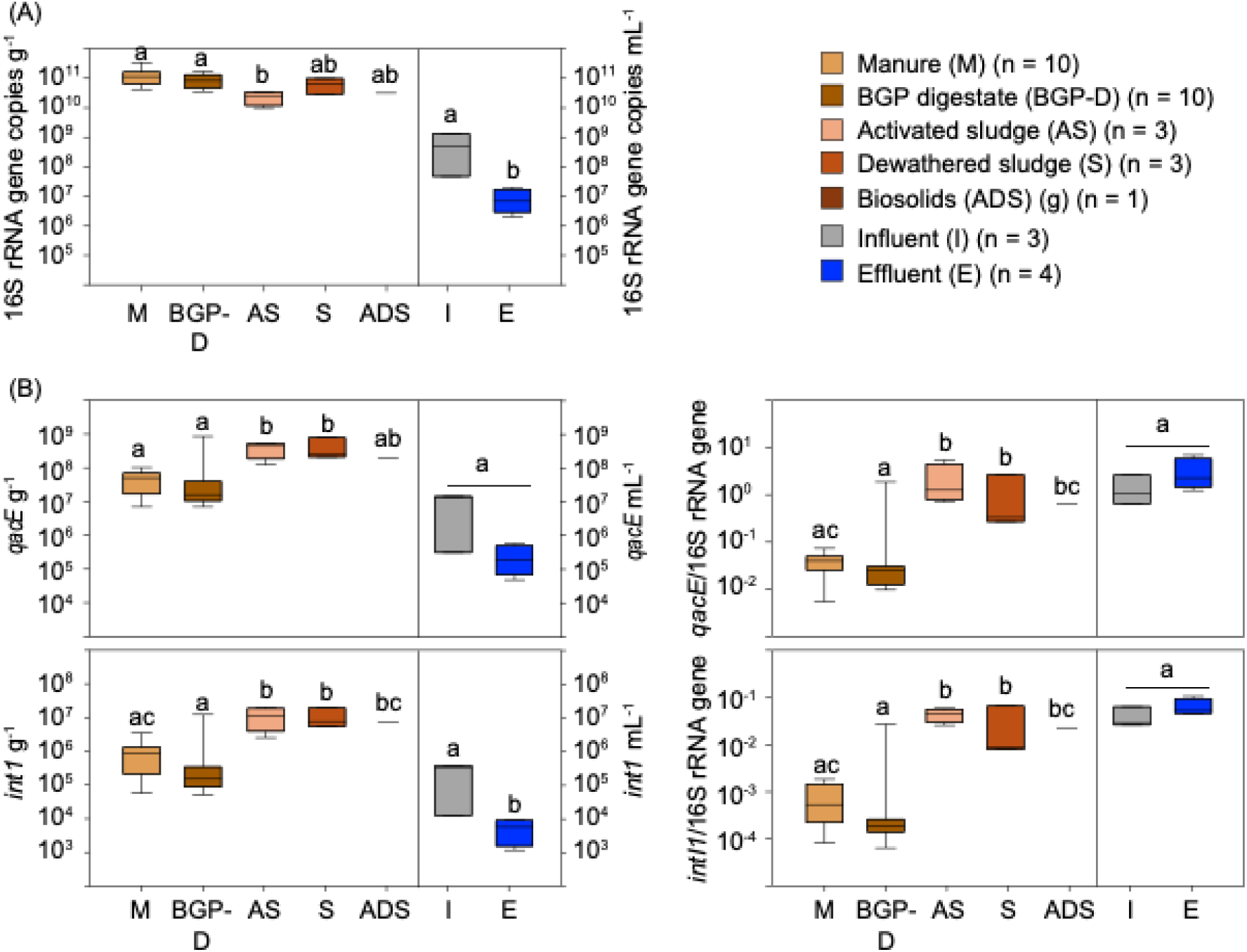
Abundance of *Bacteria,* the QAC resistant indicator genes *qacE/qacEΔ1,* and the mobile genetic element indicator gene *intI1* in livestock and human waste streams. Data based on qPCR analysis. Absolute abundances are represented as gene copy numbers per mL or g samples. A, Abundance of *Bacteria* estimated by the quantification of bacterial 16S rRNA gene copies. B, Absolute and relative abundance of *qacE/qacEΔ1* and *intI1* gene copies. Relative abundances are given as fraction of the detected 16S rRNA gene copies. Different letters at top of the bars indicate significant differences (p<= 0.05); solid samples (M, BGP-D; AS, S, ADS) were tested with Conover-Iman post-hoc comparisons using adjusted p-value; liquid samples (I, E) were tested with Wilcoxon rank-sum tests (one-sided).

### 3.3 Total and relative abundance of RGs in livestock and human waste streams

The genes *qacE*/*qacEΔ1* (QAC resistance indicator gene) and *intI1* (mobile genetic element indicator gene [79]) were detected in all studied livestock and human waste stream samples (Fig. 2B, Table S15). Absolute concentrations of both genes were in a similar range for manure and BGP digestates (most 10^6^ to 10^7^; one 10^9^ gene copies per g FW) and activated sludge, dewatered sludge, and anaerobically digested sludge samples (all in the range of 10^8^ gene copies per g FW). On a total weight basis, the abundances were significantly higher in activated and dewatered sludge samples compared to manure or BGP digestates (p_adjusted_ < 0.05; Kruskal-Wallis test with Conover-Iman post-hoc comparison, Table S17). The concentrations of *qacE*/*qacEΔ1* in the WWTP influent samples were in the range of 10^5^ to 10^7^ gene copies per mL, while the concentrations in the effluent samples were mostly 2 orders of magnitude lower compared to the influent samples. The decrease of abundance was just not significant (p = 0.03-0.05; p_adjusted_ = 0.09-0.1; one-sided Wilcoxon rank sum test, Table S19). The abundance of *intI1* showed the same trend but was one to two log scales lower in comparison to the *qacE*/*qacEΔ1* concentrations. The comparison of *qacE*/*qacEΔ1* and *int1I* genes copy numbers normalized to the bacterial 16S rRNA gene copy numbers (relative abundances), showed an identical trend for both genes: The fractions in livestock waste stream samples were significantly lower than in human waste stream samples, with the exception of the digested sludge (p_adjusted_ < 0.05; Kruskal-Wallis test with Conover-Iman post-hoc comparison; Table S20). In livestock system, no significant differences were obtained between manure and digestate samples (p_adjusted_ ≥ 0.13; Conover-Iman post hoc comparison; Table S20). In a similar manner, no differences were obtained between influent, activated sludge, effluent, dewatered or digested sludge samples (p_adjusted_ ≥ 0.13; Conover-Iman post hoc comparison; Table S20). Interestingly, the relative abundance of both genes in the effluent samples was always slightly, though not significantly, higher compared to influent samples in both, urban and rural WWTPs (p_adjusted_ ≥ 0.32; Conover-Iman post hoc comparison; Table S20).

Beside the QAC resistance and mobile genetic element indicator genes, a selection of ARGs were quantified, *sul1*, *sul2*, *tetM,* and *qnrS*, as indicator genes for sulfonamide, tetracycline, and fluorochinolone resistances (Fig. 3, Table S15). *Sul1* and *sul2* genes followed the same trend as *qacE*/*qacEΔ1* and *int1I* genes. In contrast, the absolute concentrations of *tetM* were one log scale and significantly higher in manure and digestate samples compared to activated, dewatered and digested sludge samples (p_adjusted_ < 0.05; Kruskal-Wallis test with Conover-Iman post-hoc comparison; Table S17). The relative *tetM* abundances were slightly but significantly higher for livestock systems compared to dewatered sludge (p_adjusted_ < 0.05; Kruskal-Wallis test with Conover-Iman post-hoc comparison; Table S20). Again, the relative abundance in the WWTP effluent samples was slightly higher compared to the WWTP influent samples but just not significant (p_adjusted_ = 0.11; Conover-Iman post-hoc test; Table S20). In contrast to *tetM,* the absolute and relative concentrations of *qnrS* were much higher in most of the WWTP samples (except the digested sludge) than livestock samples (p_adjusted_ < 0.05; Kruskal-Wallis test with Conover-Iman post-hoc comparison; see Tables S17, S20). The *qnrS* gene was either detected with low copy numbers or not detected (undetected in 6 out of 11 samples) in manure and BGP digestates. It was also not detected in anaerobically treated sludge of the urban WWTP. The comparison of relative abundances of *qnrS* among WWTP samples showed a higher relative abundance in influent and effluent samples compared to activated, dewatered and digested sludge but this difference was only statistically significant for both influent and effluent compared to the digested sludge (p_adjusted_ < 0.001, Conover-Iman post-hoc test; Table S20).

**Fig. 3.**
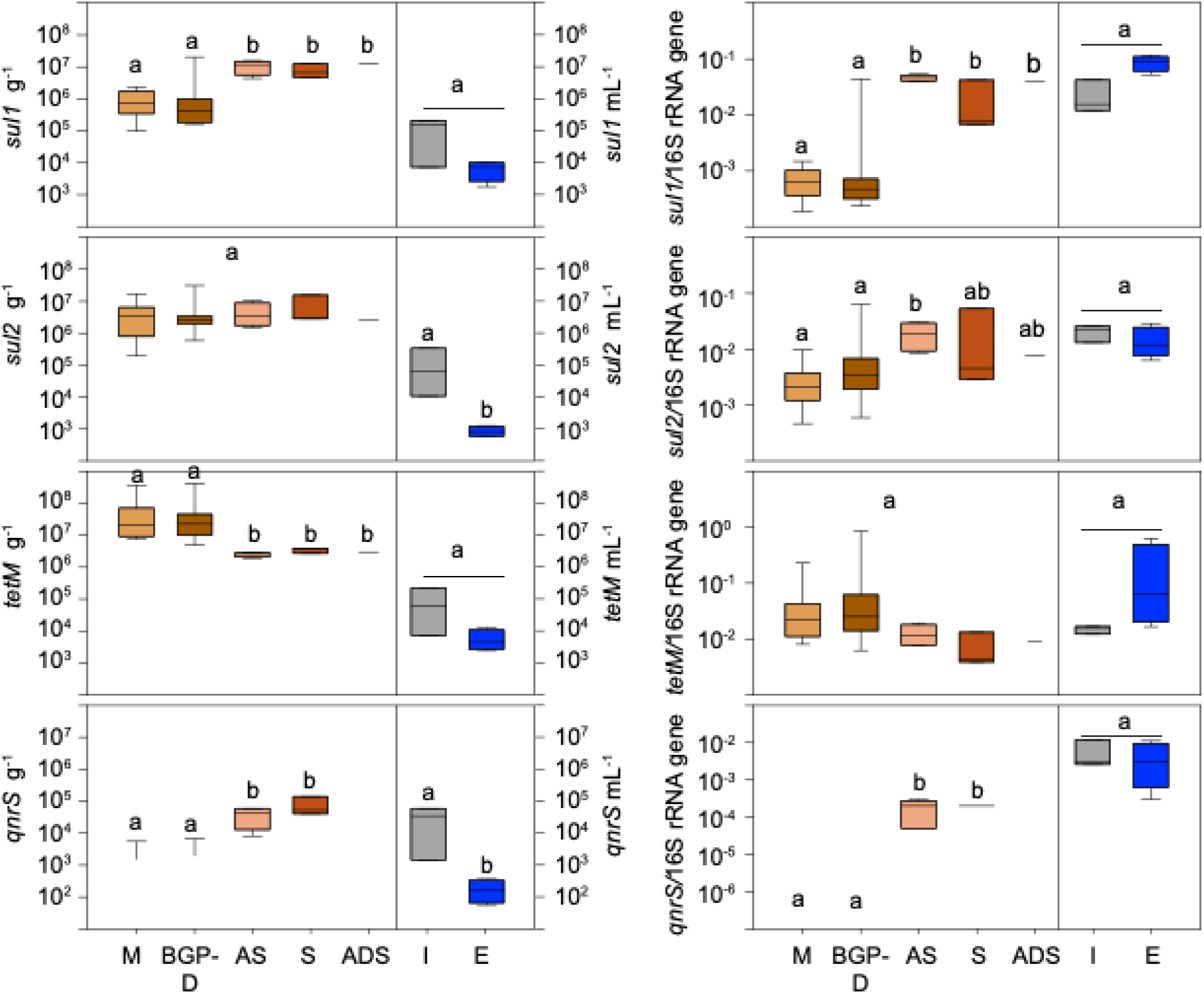
Absolute and relative abundance of antibiotic resistance genes in livestock and human waste streams. Indicator genes for sulphonamide (*sul1*/A, E and *sul2*/B, F), tetracycline (*tetM*/C, G), and ciprofloxacin (*qnrS*/D, H) resistance were tested. Data based on qPCR analysis. Absolute abundances are represented as gene copy numbers per mL or g samples. Relative abundances are given as fraction of the detected 16S rRNA gene copies (Fig. 2). Detailed figure legend (colour coding, analysed numbers of sample replicates) and statistical tests are given Fig. 2.

### 3.4 Total and BAC-C12 tolerant heterotrophic bacteria cultivated under high nutrient conditions at 37°C from livestock and human waste streams

In livestock waste streams the concentrations of heterotrophic bacteria cultivated under high nutrient conditions at 37°C (total bacteria) from manure and BGP digestates were in the range of 10^6^ and 10^8^ CFUs g^-1^ (Fig. 4A, Table S15), without significant differences (p_adjusted_ > 0.05) among sample types (manure vs. BGP digestates; Table S17) and animal source (dairy vs. pig farms; Table S16). For one out of nine manure samples and five out of nine BGP digestates, no BAC-C12 tolerant bacteria could be cultivated by the spread plating approach. For the remaining manure and BGP digestates, the concentrations of cultivated bacteria tolerant to 50 or 100 µg BAC-12 mL^-1^ were 2 to 5 log scales lower than the concentrations of bacteria cultivated without BAC-C12 under the same conditions.

**Fig. 4.**
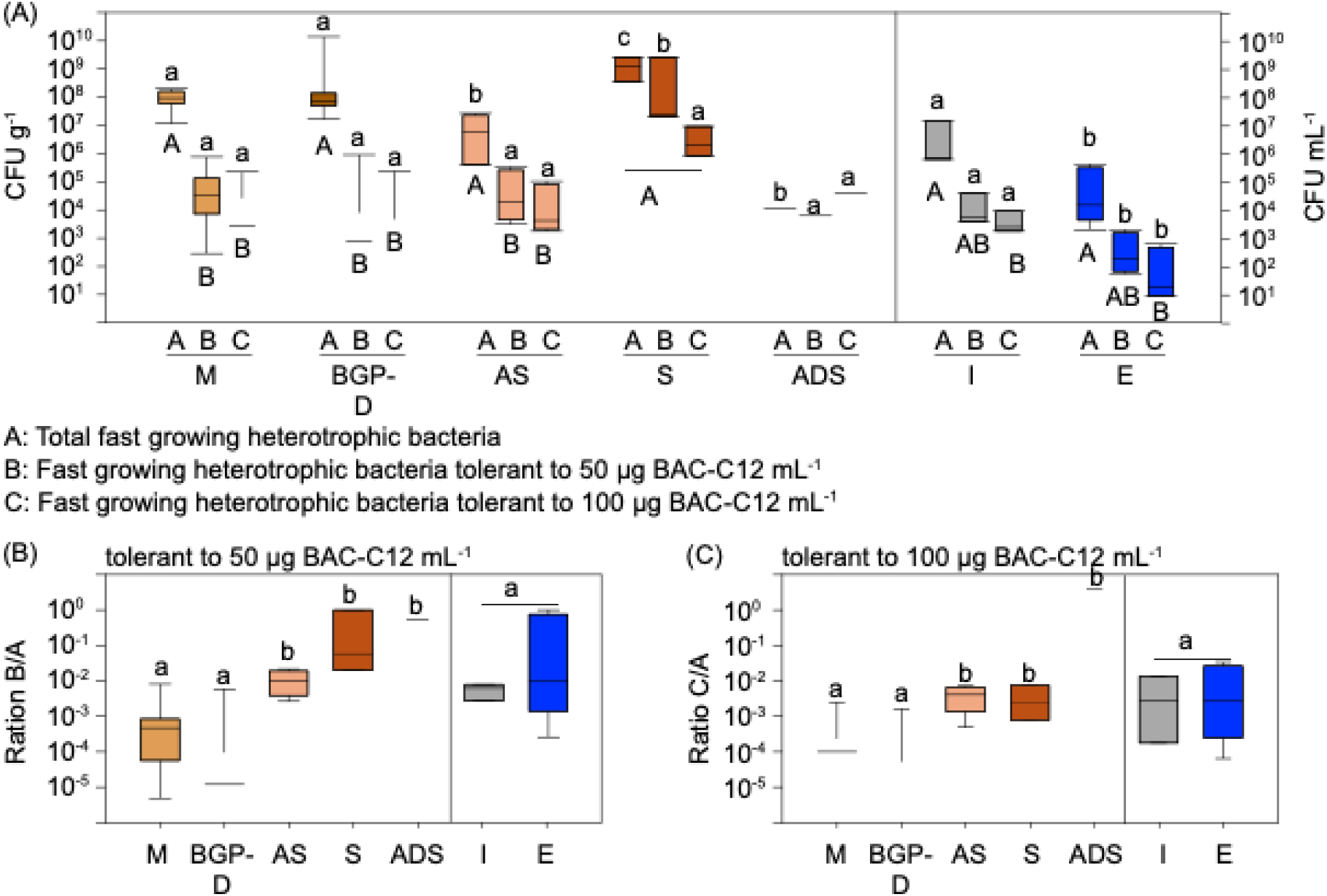
Concentrations of cultivated total fast growing heterotrophic bacteria and BAC-C12 tolerant subfractions in livestock and human waste stream samples. (A) Absolute concentrations of total cultured fast growing heterotrophic bacteria and those cultivated in the presence of 50 and 100 µg BAC-C12 mL^-1^. Concentrations are given as colony forming units (CFUs) per mL or g. (B, C) Relative abundance of BAC-C12 tolerant bacteria represented as fraction of the total cultivated bacteria. Cultivation was performed on MH agar at 37°C under oxic conditions for 48 h. Detailed figure legend (colour coding, analysed numbers of sample replicates) and statistical tests are given Fig. 2. In (A) statistical tests were done for A, B, and C, separately, according to Fig. 2. In addition, in (A), A, B, C were compared per samples type using One Way ANOVA with the Dunńs post hoc test using Bonferroni corrected p values (PAST5). Results of this test are given in addition as capital letters below bar plots.

In human waste streams, the concentrations of cultivated total bacteria in rural and urban WWTP were strongly reduced from influents (10^5^ to 10^7^ CFUs mL^-1^) to effluents (10^3^ to 10^4^ CFUs mL^-1^; p = 0.03; p_adjusted_ = 0.09; one-sided Wilcoxon rank sum test; Table S19). This corresponds to elimination rates of 97.0 to 99.7% during wastewater treatment. In the presence of 50 and 100 µg BAC-C12 mL^-1^ the concentrations in influent and effluent samples were also strongly reduced by 2-4 log scales (p = 0.03; p_adjusted_ = 0.09; one-sided Wilcoxon rank sum test; Table S19). The reduction was up to 1 log scale higher in the presence of 100 compared to 50 µg BAC-C12 mL^-1^.

The concentrations of cultivated total bacteria in activated sludge samples were in the range of 10^5^ and 10^7^ CFUs g^-1^ without remarkable differences between the rural and urban WWTPs. The concentrations of bacteria cultivated from activated sludge in the presence of 50 and 100 µg BAC-C12 mL^-1^ were 1.7 to 3.3 log scales lower than in culture without BAC-C12 addition.

Dewatered sludge samples of the rural WWTP contained concentrations of cultivated total bacteria in the range of 10^8^ to 10^9^ CFUs g^-1^. Those samples exhibited (pre and post-pandemic) the highest concentrations of bacteria cultivated in the presence of 50 and 100 µg BAC-C12 mL^-1^. Like activated sludge, the concentrations were mostly 1.7 to 3.1 log scales lower in the presence of 50 and 100 µg BAC-C12 mL^-^ than in the absence of BAC-C12. However, for the post-pandemic sample collected in March 2021, the concentration of bacteria cultivated in the in the presence of 50 µg BAC-C12 mL^-1^ was equal to the concentration of bacteria cultivated without BAC-C12.

The concentration of cultivated total bacteria from the anaerobically digested sludge of the urban WWTP collected in June 2021 was 1.1 x 10^4^ CFUs g^-1^. The concentrations of bacteria tolerant to 50 BAC-C12 mL^-1^ was only slightly lower (0.3 log scales) while the concentration of bacteria tolerant to 100 µg BAC-C12 mL^-1^ was even 0.5 log scales higher.

Expressed as relative abundances, the fractions of cultivated BAC-C12 tolerant bacteria were significantly lower for livestock compared to human waste stream samples (Fig 4B,C; p_adjusted_ < 0.05; Kruskal-Wallis test with Conover-Iman post-hoc comparison; Table S20). No significant difference in the fraction of bacteria tolerant to 50 µg BAC-C12 mL^-1^ was observed between raw and digested manure. In human waste streams, the fraction of bacteria tolerant to 50 µg BAC-C12 mL^-1^ was in the same range for influent and activated sludge and slightly, but not significantly, higher for effluent, dewatered sludge, and anaerobically digested sludge) (Fig. 4B; Table S20). More variable values were thereby obtained for waste stream output compared to input samples. For cultivated bacteria tolerant to 100 µg BAC-C12 mL^-1^ the fraction was significantly lower in manure compared to BGP digestates (p_adjusted_ = 0.04; Conover-Iman post-hoc comparison; Table S20). The fractions of BAC-C12 tolerant bacteria cultivated in the presence of 100 µg BAC-C12 mL^-1^ were in a similar range for influent, effluent, activated sludge, and dewatered sludge (rural system), but much higher for anaerobically digested sludge from the urban WWTPs (Fig. 4C).

### 3.5 Correlations between the content of QACs, RGs, and cultivated QAC tolerant bacteria, in the studied waste stream systems

Combined correlation analysis of all investigated solid samples in the two systems (i.e. manure, BGP digestates, as well as activated, dewatered and digested sludge) showed significant positive correlations between all of the following: dry weight concentrations of BAC-C12, BAC-C14, ΣBACs, ΣQACs, the relative abundance of *qacE*/*qacEΔ1*, *intI*1, *sul1* and *qnrS* RGs, as well as the fraction of BAC-C12 tolerant bacteria cultured in the presence of 50 µg BAC-C12 mL^-1^ (p_adjusted_ < 0.05, Spearman correlation; Fig. S4). The fraction of QAC tolerant bacteria cultured in the presence of 100 µg BAC-C12 mL^-1^ correlated significantly positively with the fraction cultured at 50 µg BAC-C12 mL^-1^, as well as BAC-C14, ΣBACs, ΣDADMACs, ΣQACs, *qacE*/*qacEΔ1*, *intI*1, *sul1*, and *sul2*. Only the *tetM* RG was significantly negatively correlated with BAC-12 tolerance at 50 µg BAC-C12 mL^-1^ and showed no relationship with QAC concentrations or other RG abundances.

Considering relative abundances across all investigated samples (including the wastewater), a similar pattern was observed (Fig. S5). The relative abundances of the *qacE*/*qacEΔ1*, *intI*1, *sul1*, *sul2* and *qnrS* RGs all correlated positively with each other, with the fraction of BAC-C12 tolerant bacteria cultured in the presence of 50 µg BAC-C12 mL^-1^, and, except for *qnrS*, also with the fraction of BAC-C12 tolerant bacteria cultured in the presence of 100 µg BAC-C12 mL^-1^. The strongest correlations were observed between *qacE*/*qacEΔ1*, *intI*1 and *sul1* (Spearman’s ρ >0.9). Again, only *tetM* showed a negative correlation with the fraction of BAC-C12 tolerant heterotrophic bacteria.

When looking separately at livestock systems, BAC-C14 and ΣBACs correlated positively with the relative abundance of *sul2* (Fig. 5A). The fractions of *qacE*/*qacEΔ1*, *intI*1 and *sul1* correlated with each other, and *sul2* with *tetM*. However, no significant correlations were found between the fractions of BAC-C12 tolerant bacteria and QAC concentrations or any investigated RGs.

**Fig. 5.**
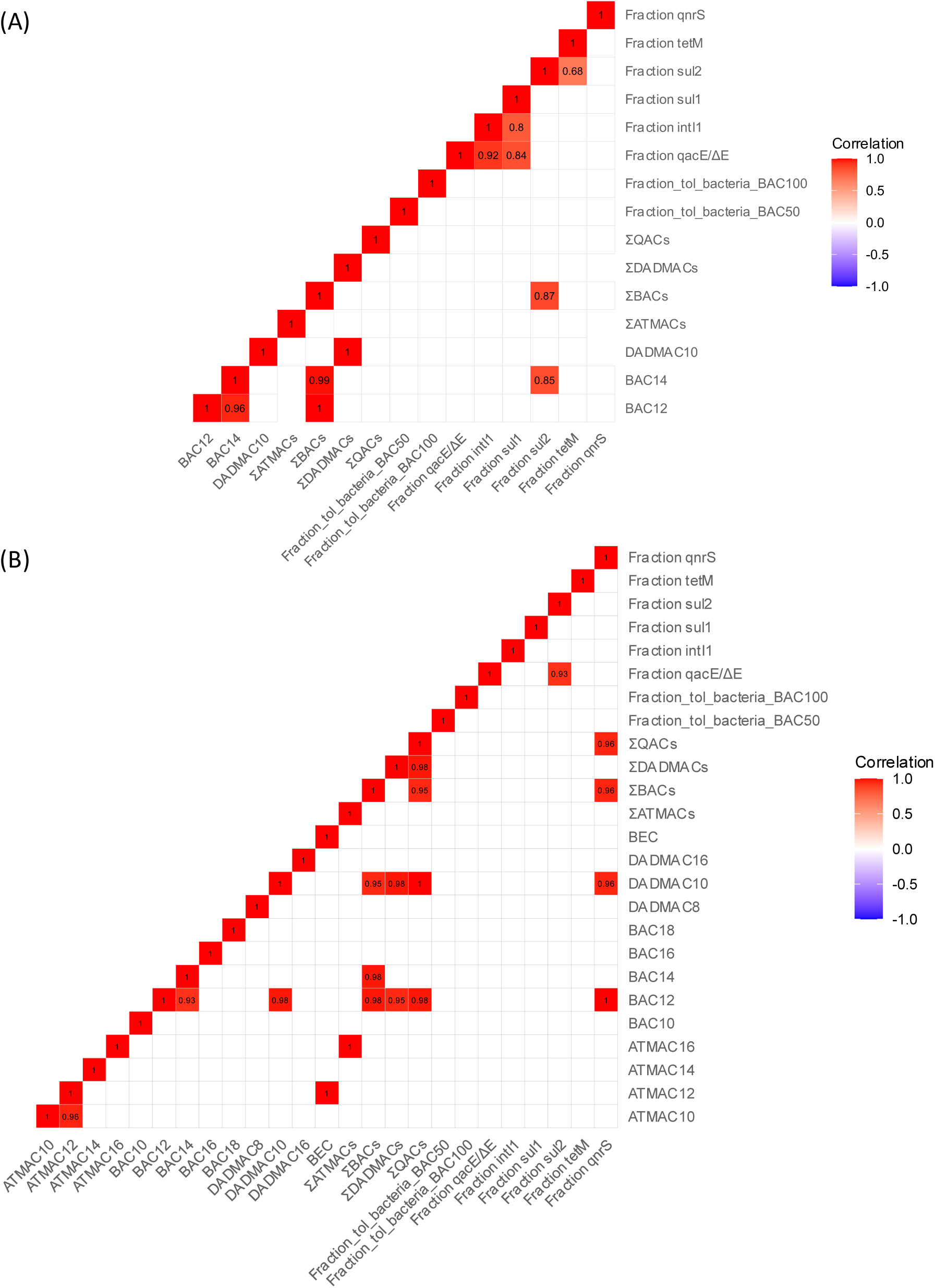
Spearman correlations between QAC concentrations, the fraction of BAC-C12-tolerant bacteria, and relative gene abundances. Only significant correlation coefficients are shown (p_adjusted_ < 0.05). Top: livestock system (manure and BGPs, n=22). Bottom: WWTP sludges (activated sludge, dewatered sludge and anaerobically digested sludge; n=8). Among wastewater samples (WWTP influent and effluent, n=7) no significant correlations were found (see Table S21).

In WWTP sludges, several QACs showed significant positive correlations with each other, while *qacE*/*qacEΔ1* correlated positively with *sul2. B*etween QACs and RGs significant positive correlations were only observed for *qnrS*, BAC-C12, DADMAC-C10, ΣBACs and ΣQACs (Fig. 5B). Similarly, in wastewater samples (Table S21) no significant correlations were found between QAC concentrations, relative abundances of BAC-C12 tolerant bacteria, or relative abundances of RGs, which may be due the small sample size (n=7).

### 3.6 Effect of the SARS-CoV-2 pandemic on QAC concentrations, QAC RGs, and cultivated QAC tolerant bacteria in the studied waste streams

In livestock farm samples, no clear effect of the SARS-CoV-2 pandemic was visible. While QACs were detected in one pre-pandemic and three post-pandemic samples in BGP-1, the opposite was true for BGP-2. After the start of the pandemic there were no significant increase in ∑QAC concentrations, RGs, or the fraction of cultivated BAC-C12 tolerant bacteria considering all aggregated samples, nor after differentiating between manure vs. digestates or individual farms (p > 0.05, one-sided Wilcoxon rank sum test; Table S22).

∑QAC concentrations in activated sludge of the rural WWTP stayed nearly identical over the three studied years 2019, 2020, and 2021. While ATMACs C14-C16, BAC-C18, and DADMAC-C16 showed tentatively higher concentrations post-pandemic, this could not be evaluated statistically. ∑QAC concentrations in dewatered sludge decreased slightly from 2019 to 2020 (−24%), followed by a strong increase in 2021. Compared to 2019 levels, ∑QACs rose by +114% (+145%∑ATMACs, +102% ∑BACs, and +115% ∑DADMACs). Along with this, a strong increase in the relative abundance of cultivated BAC-C12 tolerant bacteria was observed both in 2020 (+193%) and 2021 (+5016%), but only for the concentration of 50 µg BAC-C12 mL^-1^. In the ADS of the urban WWTP, QAC concentrations also showed a strong increase in 2021 compared to pre-pandemic levels in 2016 determined by [77]: +62% ∑ATMACs, +563% ∑BACs, + 319% ∑DADMACs and +411% ∑QACs. Data for cultivated BAC-C12 tolerant bacteria are not available for equal pre-pandemic samples. In the effluent of the rural WWTP,

∑QAC concentrations tentatively increased by 173% from 2019 to 2020 and by 162% from 2019 to 2021 while the fraction of cultivated BAC-C12 tolerant bacteria was increased by 6900% in 2020 but not 2021. However, since sampling was conducted in different months each year and individual QACs were only detected in near-LOD concentrations and at 1-2 time points in the effluent, these apparent trends should be viewed with cautions.

### 3.7 Phylogenetic identification of abundant total and BAC-C12 tolerant bacteria cultivated from livestock and human waste streams

In total, 438 bacterial strains cultivated from the waste stream systems were preserved for subsequent analysis. Among those, 171 strains were cultivated without BAC-C12 (79 from livestock and 92 from human waste streams) and 267 strains in the presence of either 50 or 100 µg BAC-C12 mL^-1^ (137 from livestock and 130 from human waste streams). The cultivated strains were assigned to 34 genera of four phyla, *Actinobacteriota, Bacteroidota*, *Bacillota*, and *Proteomonadota* (Fig. 6).

**Fig. 6.**
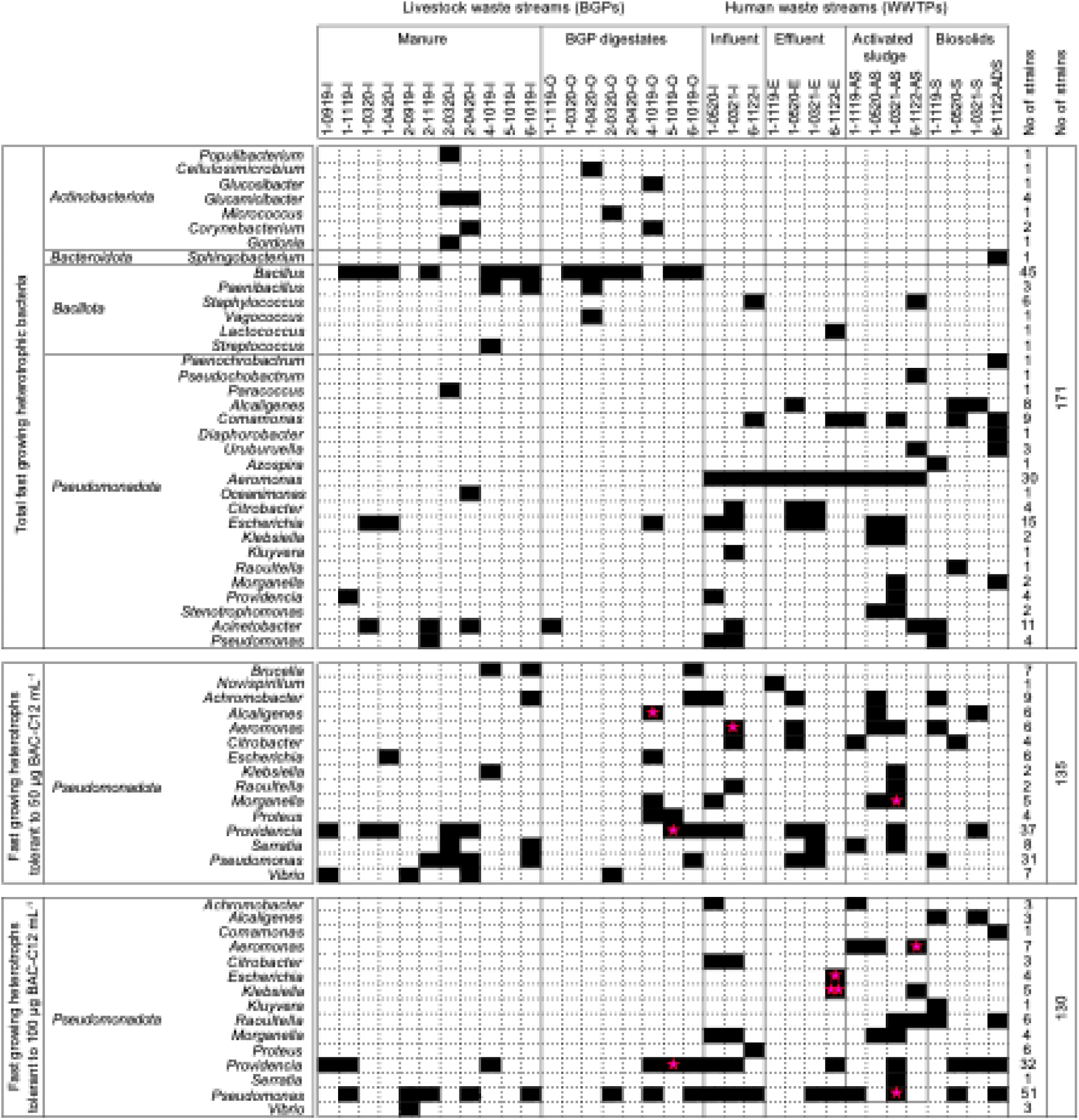
Overview of the phylogenetic placement of the potential pathogenic and BAC-C12 tolerant potential pathogenic bacteria cultivated from livestock and human waste streams. Bacteria were either cultivated under oxic conditions at 37°C on MH (171 strains) or MH supplemented with of 50 and 100 µg BAC-C12 mL^-1^ (135 and 130 strains). Pink stars indicate the presence of *qacE/ΔE* gene; number of stars indicate the number of strains that contained a *qacE/ΔE* gene.

Without BAC-C12 as selective agents, both, Gram-positive and Gram-negative bacteria, including strains of all 34 genera were cultivated. The diversity of cultivated taxa was similar in livestock and human waste streams, but the phylogenetic assignment of the cultivated strains was different between the two systems. Gram-positive bacteria were the predominantly cultivated bacteria from livestock samples (77% of livestock strains), while Gram negative bacteria were predominantly cultivated bacteria from human waste streaks (92% of the WWTPs strains). Strains of only four genera were cultivated from both systems including all sample types: *Escherichia/Shigella*, *Acinetobacter*, *Providencia*, and *Pseudomonas* (all *Pseudomonadota*). The remaining livestock strains were assigned to 13 different genera including 11 Gram-positive genera. The dominating Gram-positive genus was *Bacillus* (57% of livestock strain) cultivated from both manure and BGP digestate samples. In contrast, the dominating genus cultivated from human waste streams (from all influent, activated sludge, and effluent samples) was *Aeromonas* (33% of the WWTP strains). *Aeromonas* spp. strains were not cultivated from sludge samples. Other genera which represented at least 3 to 10% of the WWTP strains were *Comamonas*, *Alcaligenes*, *Staphylococcus*, *Kleyvera,* and *Uriburuella*. The remaining strains were assigned to further 11 less abundant genera including several *Enterobacteriales* genera, among those, *Klebsiella*, *Morganella,* and *Raoultella*.

The cultivated BAC-C12 tolerant bacteria were assigned to only 17 genera, all representing Gram-negative bacteria. *Gamma-* and *Alphaproteobacteria* were cultivated in the presence of 50 µg BAC-C12 mL^-1^, while only *Gammaproteobacteria* were cultivated in the presence of 100 µg BAC-C12 mL^-1^. In both, livestock and human waste stream samples, the main fraction of strains was assigned to the *Providencia* and *Pseudomonas*. The two genera represented together 50% and 63.8% of the strains cultivated in the presence of 50 and 100 µg BAC-C12 mL^-1^, respectively. Eight further genera were cultivated in the presence of BAC-C12 from both, livestock and human waste streams, namely *Achromobacter, Alcaligenes, Escherichia, Klebsiella, Morganella, Proteus,* and *Serratia.* In the presence of 50 µg BAC-C12 mL^-1^ those genera were cultivated from both waste streams or only livestock samples, while in the presence of 100 µg BAC-C12 mL^-1^ they were only cultivated form human waste streams. Two additional genera were cultivated only from livestock samples, *Vibrio* and *Brucella.* Both genera were cultivated in the presence of 50 µg BAC-C12 mL^-1^ and only *Vibrio* in the presence of 100 µg BAC-C12 mL^-1^. Six further genera were only cultivated from human waste streams in the presence of BAC-C12 (50 and/or 100 µg BAC-C12 mL^-1^), namely *Aeromonas, Citrobacter, Roultella, Novispirillum*, *Comamonas*, and *Kluyvera*.

### 3.8 BAC-C12 and DADMAC-C10 tolerance of cultivated bacteria – differences among livestock and human waste streams

The MICs for BAC-C12 and DADMAC-C10 were determined for 340 of the cultivated strains including 143 strains cultivated without BAC-C12 (47 Gram-positive and 96 Gram-negative strains) and 197 strains cultivated in the presence of BAC-C12 (all Gram-negative strains). For both, Gram-positive and Gram-negative bacteria, the MIC distribution patterns for both tested QACs showed a sigmoid curve with a two dilutions lower MIC peaks for Gram-positive bacteria (Fig. 7A, B). The MIC50 and MIC90 values (concentrations that inhibited 50% and 90% of the tested bacteria) for Gram-positive bacteria were 3.12 and 25 µg mL^-1^ for BAC-C12 and 1.25 and 5 µg mL^-1^ for DADMAC-C10, respectively. In contrast, the MIC50 and MIC90 values for Gram-negative bacteria were 25 and 50 µg mL^-1^ for BAC-C12 and 5 and 10 µg mL^-1^ for DADMAC-C10. Gram-negative bacteria cultivated in the presence of BAC-C12 showed a decreased susceptibility to both QACs. Again, a sigmoid curve distribution pattern was obtained which was shifted to higher MIC values (Fig. 7C). The MIC50 and MIC90 values were increased by two dilutions to 100 and 200 µg mL^-1^ for BAC-C12 and 20 and >20 µg mL^-1^ for DADMAC-C10 for those strains. The MIC distribution patterns showed no clear distinction among strains of different genera.

**Fig. 7.**
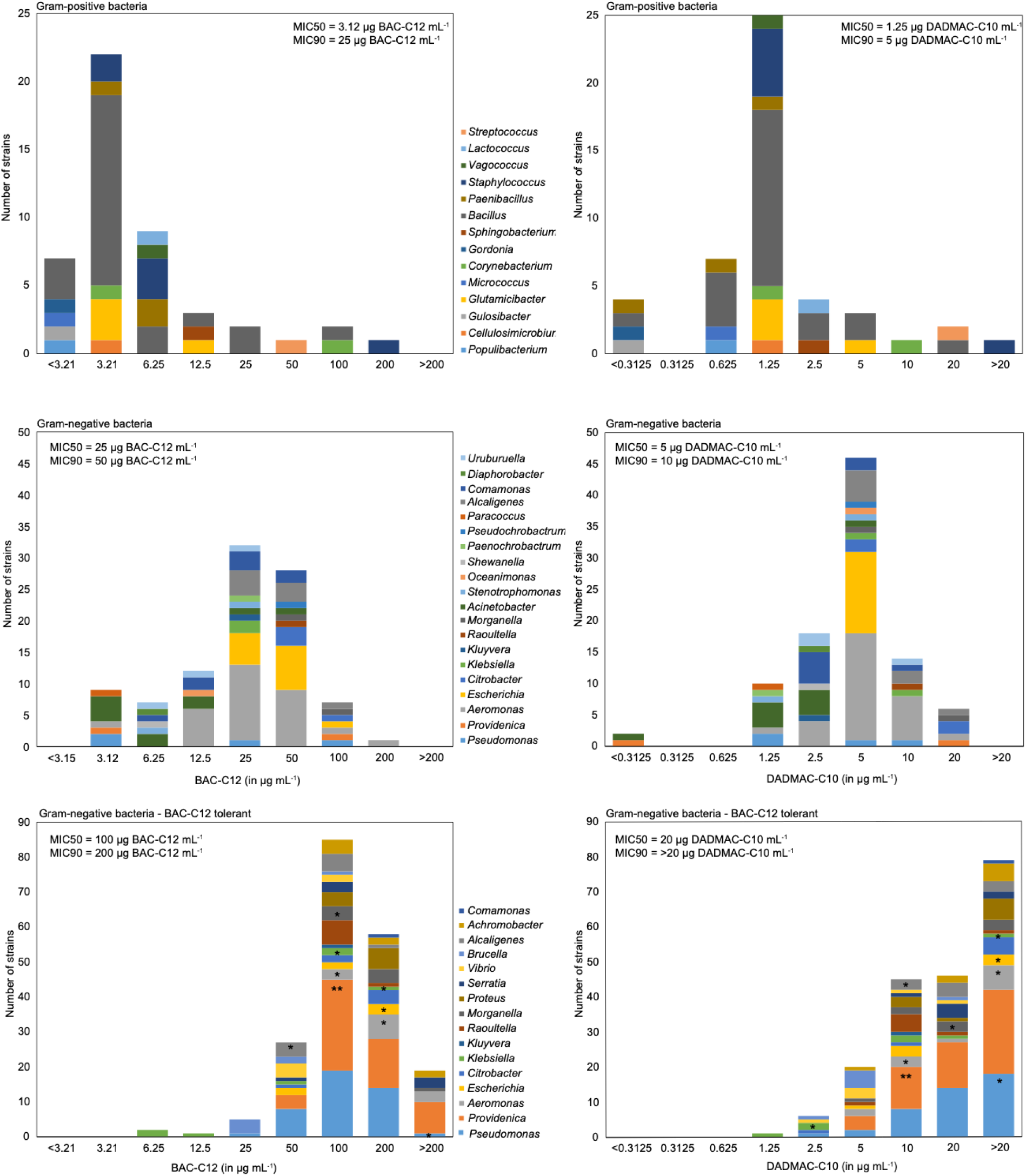
Overview of minimal inhibitory concentrations for BAC-C12 and DADMAC-C10 obtained for cultivated bacteria from livestock and human waste stream samples. Analysis was performed separately for strains cultivated under non-selective conditions (total bacteria) and those cultivated in the presence 50 and 100 µg BAC-C12 mL^-1^ (BAC-C12 tolerant bacteria). Gram-positive bacteria (n=47) and Gram-negative bacteria (n=96) cultivated under non-selective conditions and Gram-negative bacteria cultured in the presence of BAC-C12 (n=197) were separately analysed. *: indicated the presence of a strain with a *qacE/ΔE* gene which always occurred in combination with an *intI1* gene.

A separate comparative analysis was performed for strains form the three most abundant genera cultivated in the absence and presence of BAC-C12 (*Aeromonas*, *Escherichia/Shigella*, *Alcaligenes*). The analysis showed a clear MIC shift with increased MIC50 and MIC90 values at the genus level for both tested QACs if the bacteria were cultivated in the presence of BAC-C12 (Fig. S6). We included also the genera *Providencia* and *Pseudomonas* in this comparative test, but only very few strains of these genera were cultivated in the absence of BAC-C12. However, also for those few strains a shift to strains with higher QAC MICs was observed when the bacteria were cultivated in the presence of BAC-C12 (Fig. S6).

A genus level-based comparison of the MIC distribution patterns of QAC tolerant strains from livestock and human waste streams was performed for two genera representing most QAC tolerant strains from both systems. A total of *53 Providencia* and 43 *Pseudomonas* strains were cultivated in the presence of BAC-C12 from the two waste stream systems. For both genera strains from human waste streams had higher BAC-C12 and DADMAC-C10 MIC values than those cultured from livestock waste streams (Fig. 8A). The MIC50 values of strains from human waste streams were always one to two dilutions higher than MIC50 values of strains from livestock waste streams. The comparison of this dataset per sample type showed no clear differences among manure and BGP digestates in livestock systems or between influent, effluent, activated sludge, dewatered sludge, or anaerobically digested sludge in human waste stream systems (Fig. S7A-D).

**Fig. 8.**
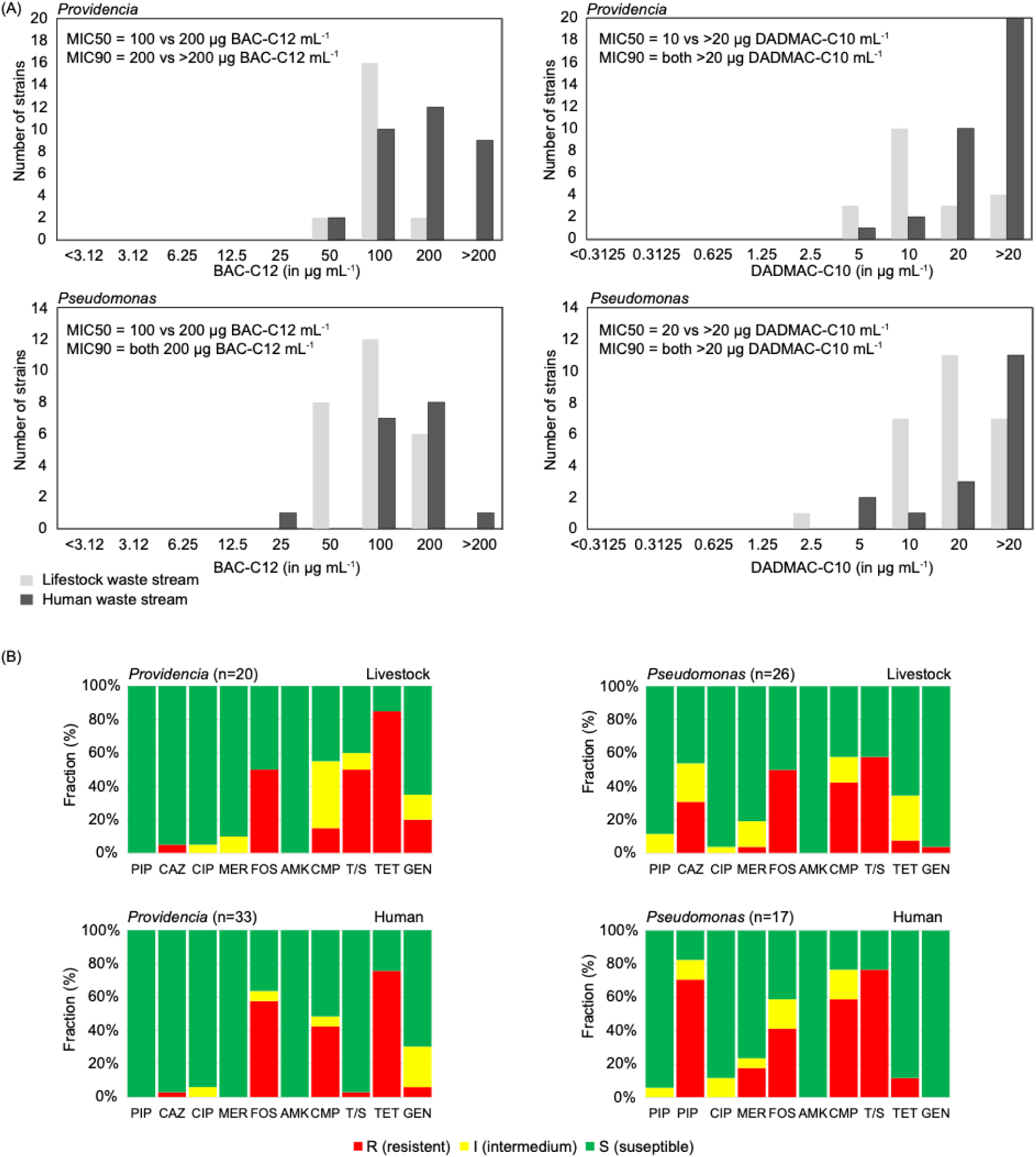
Comparison of the QAC tolerance (A) and antimicrobial susceptibility (B) of *Providencia* and *Pseudomonas* strains cultivated from livestock and human waste streams as BAC-C12 tolerant bacteria.

### 3.9 Antimicrobial resistance profiles and MDR status of cultivated bacteria

The cultivated BAC-C12 tolerant Gram-negative bacteria were subjected to AST. In total 197 strains of 16 genera were tested including the 53 *Providencia* and 43 *Pseudomonas* strains (Table S23). Antibiotics of ten different classes were tested. The classification into S and R was performed with the same cut-off values for all tested strains and all antibiotics were included in the calculation of the MAR indexes. Of the 197 tested strains 10% were resistant to piperacillin, 37% to a cephalosporine of the 3^rd^ generation (CTX or CAZ), only two and five percent were resistant to ciprofloxacin and meropenem, 52% to fosfomycin, 4% to amikacin, 37% to chloramphenicol, 30% to TMP/SMX, 44% to tetracycline, and 7% to gentamycin. Considering the MDR status of the QAC tolerant strains, 16.8% of the strains were susceptible to all tested antibiotics, 26.9% showed a resistance to one of the tested antibiotics, 14.7% to two, 25.4% to three, 11.7% to four, and 4.6% to five of the tested ten antibiotics. Again, *Providencia* and *Pseudomonas* strains were selected to compare the susceptibility profiles of strains derived from the two studied systems (Fig. 8B). No distinct differences were obtained, except a higher fraction of TMP/SMX resistant *Providencia* strains in livestock. While 50% of the livestock strains were resistant to TMP/SMX only 3% of the *Providencia* strains from human wastewaters were resistant against TMP/SMX.

### 3.10 Detection of *qacE*/*qacEΔ1* and *intI1* genes in cultivated BAC-C12 tolerant bacterial strains

Strains cultivated in the presence of BAC-C12 were screened for *qacE*/*qacEΔ1* and *intI1* (Figs. 6, 7; Table S23). Out of the 197 screened strains 10 strains (5% of the tested strains) carried the *qacE*/*qacEΔ1* and the *intI1* gene and additional 13 strains carried only the *intI1* gene (7%). None of the strains carried only *qacE*/*qacEΔ1*. All strains which carried *qacE/ qacEΔ1* and/or *intI1* were identified as *Proteobacteria*. Three of the *qacE*/*qacEΔ1* carrying strains were cultivated from digestates of the two swine manure processing BGPs, one *Alcaligenes* strain (BGP4) and two genetically identical *Providencia* strains (BGP6). Three *qacE*/*qacEΔ1* carrying strains were cultivated from the rural WWTP, one *Aeromonas* strain was cultivated from WWTP influent and one *Pseudomonas* and one *Morganella* strain from the same activated sludge sample. The remaining strains were cultivated from the urban WWTP, one *Aeromonas* strain from activated sludge, one *E. coli* and two genetically different *Klebsiella* strains from WWTP effluents. The QAC MIC values of *intI1* and *qacE*/*qacEΔ1* carrying strains were widespread over the complete detected MIC-range from 50 to >200 µg BAC-C12 mL^-1^ and 5 to >20 µg DADMAC-C10 mL^-1^, respectively. The same was obtained for the MDR status, ranging between a MAR index of 0.1 to 0.5 (Table S23). A MAR index above 0.3 was only obtained for two *qacE/qacEΔ1* carrying *Providencia* strains which were resistant to fosfomycin, amikacin, chloramphenicol, TMP/SMX, tetracycline, and gentamycin.

## 4 Discussion

Livestock and human waste streams are considered as the two main hotspots for the release of AMR into the environment [95,96]. However, considering the increasingly discussed role of biocidal compounds, especially QACs, on the transmission of AMR [12], there is a lack of investigation of the spread of bacterial QAC tolerance in those systems. This study contributes to a deeper understanding of the presence of QACs, QAC resistance genes, and QAC tolerant bacteria in both waste stream systems.

### 4.1 Occurrence of different QAC homologues in livestock and human waste streams

BAC-C12 and DADMAC-C10 were the most abundant QACs in both livestock and human waste streams. This confirms prior studies reporting their occurrence, and often high concentrations, in manure, BGP digestates, wastewater and sludge [12,26,60,97,98], which can be attributed to their widespread use in disinfectants, biocides and cleaning products [12,13,59]. BAC-C12 and DADMAC-C10 were therefore the most relevant QACs to test bacterial tolerance in this study.

Overall, a more diverse set of QACs occurred in samples from the human waste stream, reflecting diverse applications in consumer products, industry and healthcare. Similar to prior studies [47,67], ATMACs occurred in lower concentrations than BACs or DADMACs in all samples but the WWTP effluent. This may be caused by stronger sorption of QACs with increasing hydrophobicity [19,39,45], favouring particle partitioning and potentially reducing the (bio-)degradation of benzyl-substituted, long-and double-chained QACs. Relatively high fractions of short-chained ATMACs in the effluent could be explained by their hydrophilicity and the very low effluent particle content. While other studies have reported high concentrations of long-chain BACs and DADMACs in BGP digestates [97], human wastewater [99] and sewage sludge [43], the SPE methods employed in this study resulted in lower recovery rates for the most hydrophobic QACs. Consequently, some concentrations, e.g. for DADMAC-C18, may be considerably underestimated. Prior studies have also reported high concentrations of long-chained ATMACs C18-C22, DADMACs C20-22 as well as unequal-chain DADMACs in sludge [43,46], wastewater-impacted suspended particulate matter [51,100] and sediment [48,53–55]. These hydrophobic QAC homologues are also expected to contribute to the solid-bound QACs in German WWTPs, and potentially in livestock waste streams.

### 4.2 Comparative QAC emissions from human and livestock waste streams

The measured QAC concentrations generally fell within the range reported in literature for sludge (µg g^-1^) and WWTP influent (µg L^-1^) [11–13]. Overall higher concentrations in the urban compared to rural WWTP can be attributed to the presence of several hospitals and clinics in the urban catchment as Wolter et al. [26] previously observed elevated QAC concentrations in biosolids of German WWTPs receiving hospital wastewater. While wastewater particles are sometimes neglected in monitoring studies, here 76-99% of QACs in WWTP and influent and activated sludge were particle-associated. These findings agree well with studies from Sweden, Denmark and France reporting approximately 90% of incoming QACs in the solid wastewater phase [9,45,101], and underpin the importance of accounting for particle partitioning in mass balance and fate studies on QACs. Moreover, as Mongelli et al. [45] demonstrated that QACs bound to AS are still biodegradable and thus accessible to microorganisms, this new study indicates that particle-associated microorganisms may play a crucial role in the transmission of QAC tolerance and AMR from WWTPs.

Data on QACs in livestock waste is still very scarce. Buijs et al. [60] reported concentrations of BACs C12-C14 and DADMAC-C10 up to hundreds ng/g dw in liquid manure from Dutch farms while an Austrian study found ∑QAC levels (ATMACs C12-C16, BACs C12-C18, DADMACs C10-C18) between 350 ng g^-1^ and 180 µg g^-1^ in BGP digestates [97,102]. Moreover, Larsson et al. [98] reported concentrations of 800 µg BAC-C12 L^-1^ and 170 µg BAC-C14 L^-1^ in digestates of Danish BGPs processing >75% swine manure. At the upper end, these concentrations are much higher than concentrations in the current study. However, the Austrian study included BGPs processing both animal and human waste (e.g. from catering, households, and food industry) and reported highest concentrations for plants using slaughter and food waste as input. In contrast, QAC concentrations in the Austrian BGP digestates originating mainly from manure fell in a similar range as peak concentrations measured in pig manure digestates in this study. Overall higher QAC concentrations occurred in pig compared to cattle manure, which is in line with the excessive concentrations reported by Larsson et al. [98] in swine manure digestates and can be explained by stricter disinfecting protocols in pig farming [41,57,58].

Overall, biofertilizers from the human waste stream can thus be expected to contain 10-100-fold higher QAC concentrations than raw or anaerobically treated livestock manure. In Germany, about 120 Mt raw liquid manure and 0.3 Mt sewage sludge are annually applied on farmland [92,94], making manure by far the most widely used biofertilizer. Nevertheless, based on our data, the estimated QAC emission loads into German agricultural soils are still up to 10-fold higher from digested sewage sludge (13 t/a) compared to manure (1.4 t/a) or BGP digestates (2 t/a) and direct discharge with WWTP effluent (<1 t/a). These figures may shift in the future since the German Sewage Sludge Ordinance [103] prohibits the land application of sewage sludge from WWTP with more than 50,000 person equivalents starting from 2032. Already, the proportion of sludge disposed of on agricultural land has been decreasing from approx. 30% in 2010 to 13% in 2023 [92,104]. While this could achieve a reduction of nationwide emissions of sludge-borne pollutants, it may not necessarily decrease inputs into individual fields as QAC concentrations in German biosolids were not directly affected by WWTP size [26] and QACs could still reach fields with sludge from small WWTPs. Compared to Germany, in countries where agricultural sludge application is more common, e.g. France, Portugal, Spain, Ireland, the UK, Sweden and Denmark [105,106], sludge-associated QAC release into soils may be even higher than in Germany. Overall, environmental QAC emission from WWTP biosolids are therefore expected to exceed release via livestock manure and discharge with WWTP effluent. Manure and wastewater from swine farms may however also play an important role in QAC, pathogen and AMR release [58]. Further research should investigate the fate and microbial effects of QACs entering soil via manure or biosolids amendment as organic fertilizers, and via irrigation with reclaimed wastewater.

### 4.3 Potential of WWTPs and BGPs to reduce QAC emissions

In WWTPs, prior studies observed high QAC removal rates from the wastewater of approximately 90 to >99% [38,42,47,99], which is in line with findings from our study. While QACs are biodegradable under oxic conditions [107–109], they also sorb strongly to activated sludge solids [38], which may slow down degradation [110]. Hence, a large fraction of QACs is transferred to dewatered or digested sludge. According to our and prior mass balance studies [42] [38], roughly 10-70% of incoming QACs are transferred to sludge rather than eliminated. Thus, WWTP can significantly reduce QAC concentrations in the “liquid waste stream” and cut direct emissions into surface waters or soils irrigated with treated wastewater. However, when looking at the “solid waste stream”, in this study neither AS treatment nor anaerobic manure digestion substantially reduced QAC concentrations in solid waste products. Large amounts of QACs may therefore still enter agricultural soils sorbed to biosolids or manure. Exploring the fate and effects of biofertilizer-borne QACs thus remains an important direction for future research. Moreover, it is still unclear whether constant low-level discharge with the treated effluent may also exert selective pressure for microbial adaptation and the spread of AMR in surface waters, especially given the high relative abundance of QAC-tolerant bacteria.

### 4.4 Abundance of the QAC resistance indicator genes in waste stream systems and cultivated QAC tolerant bacteria

The indicator genes for QAC resistance *(qacE/qacEΔ1)* together with the indicator gene for anthropogenic pollution (*intI1*) were detected in all livestock and human waste stream samples with similar occurrence patterns. A consistent detection of *qacE/qacEΔ1* was also reported for manure of other German livestock farms [111,112]. Wolters et al. [111] found a significant reduction of the relative abundance of *qacE/qacEΔ1*, *intI1*, and ARGs during the anaerobic digestion in BGPs. We obtained neither for thermophilic nor mesophilic BGP processing a reduction of *qacE/qacEΔ1, intI1,* or tested ARGs. This was also observed during lab scale anaerobic digestion of sludge [11].

In human waste streams we detected *qacE/qacEΔ1*, *intI1,* and ARGs (*sul1*, *sul2,* and *tetM*) with a similar relative abundance in non-digested dewatered sludge and anaerobically digested sludge (biosolids). However, elimination rates could not be determined because samples were obtained from different WWTPs. The *qacE/qacEΔ1* and ARG concentrations in biosolids were similar to those detected in biosolids of other German WWTPs [26]. The biosolid sample studied here (WWTP6-ADS) contained the highest QAC concentrations. However, Wolters et al. [26] found that the occurrence of *qacE/qacEΔ1* genes does not correlate with QACs concentrations. In other studies, it was shown that the relative abundance of *qacE/qacEΔ1* increased in activated sludge enriched with QACs [113] indicating a correlation of QAC and *qacE/qacEΔ1* abundances. While most of the tested ARGs showed similar absolute and relative abundances as *qacE/qacEΔ1* in the tested samples, *qnrS* was only well detectable in liquid waste streams samples including influent, effluent, and activated sludge. This may be due to the predominant occurrence of *qnrS* carrying bacteria in a free-living lifestyle.

The direct comparison of the relative abundance of *qacE/qacEΔ1* and *intI1* in organic fertilizers from livestock waste streams (manure, BGP digestates) and human waste streams (biosolids) is an important issue of risk assessment. The introduction of *qacE/qacEΔ1* into the agricultural system is much higher by the application of biosolids compared to manure and BGP digestates if relative abundances are compared. As shown by Wolters et al. [26] the abundance of *qacE/qacEΔ1* in biosolids is independent to the size of the WWTP. Gaze et al. [114] showed that *qacE/qacEΔ1* introduced with organic fertilizers into soil can be still detected up to two years after fertilization. Beside organic fertilizers, irrigation with wastewater also contributes to the accumulation of *qacE/qacEΔ1* genes in soils. This was shown for agricultural fields in Mexico long-term irrigated with untreated wastewater [115,116]. Depending on the regulations in different countries, treated and partially untreated wastewater is used for irrigation of agricultural fields [117]. In Germany, effluent water of WWTPs is released into rivers adjacent to the WWTPs but with increasing water demand field application is under discussion. We detected *qacE/qacEΔ1* consistently in effluent waters of the two studied WWTPs. A consistent detection of *qacE/qacEΔ1* in treated wastewater was also reported in other studies [118,119]. Tavares et al. [120] showed that the abundance of *qacEΔ1* and *intI1* are good indicators for the presence of ARGs in WWTP influent and effluent samples. As shown by Luo et al. [119] we obtained a reduction of the absolute abundance of *qacE/qacEΔ1* in effluent compared to influent wastewater, but the relative abundance of *qacE/qacEΔ1* as well as *intI1* was higher in effluent waters. This indicates an increased abundance of *qacE/qacEΔ1* and integron carrying bacteria in effluent water bacterial community compared to the influent. The increased relative abundance of those RGs can be explained by the fact that bacterial communities of effluent water represent a mixture of activated sludge and influent microbial communities [121,122]. The circulation of activated sludge microbial communities with the return sludge for a long period with consistent exposure to disinfectants, heavy metals, and antibiotic residues may have increased the horizontal gene transfer among activated sludge bacteria. A fraction of activated sludge bacteria is then released via effluent waters into the environment.

The relative abundance of *qacE/qacEΔ1* was between 1.2 and 2.3 log_10_ scales higher than the relative abundance of *intI1* and *sul1* in all samples of both waste stream systems. The same trend was reported for manure and BGP digestates by Wolters et al. [111]. Gaze et al. [114] even reported three times higher prevalence of *qacE/qacEΔ1* compared to *intI1* in pig slurry and Jechalke et al. [115] found the same for long-term wastewater irrigated Mexican soils. The presence of complex integrons carrying two *qacE/qacEΔ1* genes or the presence of other plasmids where *qacE/qacEΔ1* occurs in another genetic context without co-occurrence of *intI1* and *sul1* can explain the differences [111,114]. While we observed significant differences for the relative abundances of *qacE/qacEΔ1* compared to *int1I* and *sul1* in livestock samples, the difference in relative abundance of *qacE/* compared to *intI1* and *sul1* was much higher in human waste streams. This indicates differences in the genetic structure of antibiotic resistant bacteria (ARBs). In a metagenomic study of effluents from urban WWTPs in Tokio, Sekizuka et al. [118] found a predominant occurrence of *sul1* followed by *qacE*. The authors concluded the predominance of *sul1* and *qacEΔ1* containing class 1 integrons as mobile genetic elements in the effluent waters. Differences in the relative abundances of *qacE/qacEΔ1, sul1,* and *intI,* among different samples indicate a different genetic structure of resistance gene carrying mobile genetic elements of *qacE* carrying *bacteria*.

We cultivated QAC tolerant bacterial strains from both waste stream systems which always carried *qacE/qacEΔ1* in combination with *intI1* (5.1% of cultivated QAC tolerant strains). In addition, BAC-C12 tolerant stains were cultivated which carried only *int1I* (6.6% of QAC tolerant strains). The overall incidence of integron positive cultivated strains was slightly higher in our study (11.7%) compared to previous studies (8.0% [31]), but the fraction of *qacE/qacEΔ1* carrying strains was lower (43.5% of the *int1I* positive strains in our study and 95.7% in the study of Gaze et al. [31]. Most of the *qacE/qacEΔ1* carrying strains cultivated in our study came from human waste streams, again confirming the correlation of QAC RG occurrence and QAC concentrations. This is in line with the observation of Gaze et al. [31], who found *qacE/qacEΔ1* carrying strains only in reed bed samples exposed to QACs. The strains that carried *qacE/qacEΔ1* and *int1I* genes in our study were assigned to the genera *Alcaligenes* and *Providencia* (livestock strains), as well as *Aeromonas, Pseudomonas, Morganella, E. coli,* and *Klebsiella* (human waste streams). No differentiation of *qacE* and *qacEΔ1* genes was performed and the abundance of further RGs and the genetic context of *qacE/qacEΔ1* in the genome of the strains was not yet studied. However, the parallel detection of *qacE/qacEΔ1* and *intI1* is at least an indication that the strains may carry class 1 integrons, which are central players in the global problem of AMR [123]. *Pseudomonas* and *Aeromonas* strains carrying *qacE/qacEΔ1* and *intI1* were also cultivated by Gaze et al. [31] from QAC contaminated reed beds. Different class1 integrons with *intI1* at the 5’CS and *qacEΔ1-sul1* fusion genes at the 3’CS were detected in wastewater derived strains of the genera *Aeromonas*, *Morganella*, *Comamonas*, *Providencia*, *Proteus*, and *E. coli* by Guo et al. [124]. All integron carrying strains analysed by Guo et al. [124] showed high resistance to several antibiotics and contained additional ARGs in the integron cassette integrated between *intI1* and *qacEΔ1.* The presence of the same integron 1 gene in strains of different genera is an indication for HGT during wastewater treatment. A similar conclusion was drawn by Kim et al. [125] who determined another gene cluster containing several efflux pump genes (RND, *sugE*, ABC transporter) in strains of different genera including *Pseudomonas*, *Achromobacter*, *Samonella*, *Citrobacter*, and *Klebsiella,* which were all cultivated from long-term BAC amended bioreactor cultures originating form river sediments. Chen et al. [126] characterized *qacE* carring *Providencia* strains of the species *P. stuartii* and *P. rettergii* which were cultivated from pig faeces. They determined carbapenem RG in the class 1 intregon cassette of those strains. Here, a broad variety of phylogenetically (phylotyping) and genetical (genotyping) different *Providencia* spp. strains were cultivated including strains closest to *P. stuartii* and *P. rettergii*. In contrast to the strains cultivated by Chen et al. [126], none of our strains carried carbapenem resistances, but a risk for RG transmission is indicated. Mbelle et al. [127] characterized a *M. morganii* strain with a new class 1 intergon. The strain was identified as an opportunistic pathogen linked to a urinary tract infection. Those examples underline the risk for human health linked to wastewater associated bacteria released into the environment.

We did not obtain higher QAC MIC values for *qacE/qacEΔ1* positive strains compared to other strains of the same genera cultivated in the presence of BAC-C12. This indicates that further resistance mechanisms must be present in other Gram-negative strains cultivated in the presence of BAC-C12. So far, little is known about the QAC resistance mechanism of environmental strains cultivated with QAC as selective cultivation supplement. More detailed genetic analysis of cultivated QAC tolerant strains is therefore required.

### 4.5 Characterization of cultivated BAC-C12 tolerant fast growing heterotrophic bacteria

There are only few studies available which performed comparable cultivation studies from environmental samples. Gaze et al. [31] cultivated QAC tolerant bacteria from wastewater contaminated reed beds and Guo et al. [128] studied soil and produce water from shallow gas production sides. Some additional studies performed first mesocosm or fermenter experiments with QAC amendments and cultivated QAC tolerant bacteria from those lab systems inoculated with natural bacteria communities of contaminated sediments [32], kitchen drain biofilms [129], sewage, soil, or activated sludge [130]. Different media, different incubation temperatures, and different QACs in different concentrations were applied which did not enable a direct comparability of the studies. A comparative overview of the cultivation conditions and cultured QAC tolerant bacterial strains is given in supplementary Table S15.

We selected BAC-C12 as selective agent because it was one of the most abundant QACs in both studied waste stream systems. MH agar incubated at 37°C was selected to focus on bacteria culturable at 37°C under high nutrient conditions because those conditions favour the growth of potential human pathogens. The detection of BAC-C12 tolerant bacteria in the two systems followed the abundance of QACs. In livestock waste stream samples containing lower concentrations of QACs than human waste stream samples, lower fractions and more samples without detectable BAC-C12 tolerant bacteria were obtained. Especially in BGP digestates the detection frequency was reduced, but overall, BAC-C12 tolerant strains could be cultivated from all different types of samples. All cultivated BAC-C12 tolerant strains of our study (all Gram negatives) also showed high MIC values for DADMAC-C10. MIC values differed depending on the structure of the tested QACs, e.g. one log10 unit lower MIC values were obtained for DADMAC-C10 compared to BAC-C12 [68].

This unique dataset of bacteria cultivated under non-selective conditions and in the presence of BAC-C12 gave some new insights into QAC tolerant bacteria in the two studied waste stream system. Several taxa which were culturable in the presence of 100 µg BAC-C12 mL^-1^ from human waste stream samples were only culturable in the presence of 50 µg BAC-C12 mL^-1^, but not 100 µg mL^-1^, from livestock samples, namely *Achromobacter*, *Serratia*, *Alcaligenes*, *Morganella*, *Klebsiella*, *Escherichia*, and *Proteus.* This is a clear indication for higher QAC tolerance of bacterial strains of those genera in the human waste stream, which is also showing a higher QAC load. Several of those genera were also detected in the other studies which cultivated QAC tolerant bacteria*. Serratia* and *Aeromonas* strains were also cultivated from QAC contaminated reedbeds and/or river sediments [31] and Guo et al. [128] cultivated *Achromobacter*, *Serratia*, and *Klebsiella* from produced water of shale gas production sites. Ertekin et al. [130] cultivated *Achromobacter, Serratia,* and *Comamonas* from BAC enrichment cultures based on sewage, activated sludge, soil, and sea sediment and Forbes et al. [129] also cultivated *Achromobacter, Alcaligenes,* and *Aeromonas* from QAC exposed kitchen drain biofilms.

Independent of the waste stream system, strains cultivated in the presence of BAC-C12 showed higher MIC values for DADMAC-C10 compared to strains cultivated on MH without QACs. MIC values of strains from both waste stream systems merged into one distribution showing a bimodal pattern, illustrating the separation of the QAC tolerant strains from the QAC susceptible population. This is clear evidence for acquired QAC tolerance in the waste stream systems. For three different bacterial genera for which enough strains were cultured from non-selective and selective conditions, namely *Aeromonas*, *Escherichia/Shigella,* and *Alcaligenes*, this observation could be confirmed at the genus level. Hausherr et al. [131] showed a similar bimodal BAC-MIC distribution pattern for *E. coli* strains cultivated from calves of fattening farms in Switzerland under non-selective cultivation conditions and gave a similar conclusion of an indication for acquired QAC resistance.

*Pseudomonas* and *Providencia* were the most abundant genera cultivated in the presence of BAC-C12 in our study from both, livestock and human waste stream systems. The large numbers of strains of those genera from both waste stream systems enabled us to compare system specific differences in more detail. Higher QAC tolerance was obtained for strains cultivated from human waste streams which also contained higher QAC concentrations compared to the livestock waste stream. In contrast we did not observe significant difference in the MDR status of *Pseudomonas* and *Providencia* from the two different waste streams except for TMP/SMX. Regarding the TMP/SMX resistance, we obtained a higher resistant rate in livestock systems for strains assigned to the genus *Providencia*.

*Pseudomonas* spp. were also the most abundant QAC tolerant bacteria in all other studies that cultivated QAC tolerant bacteria [31,128–130]. *Pseudomonas* spp. are well known for their QAC tolerance especially to BACs [132,133]. The QAC tolerance of *Pseudomonas* spp. is linked to a broad range of mechanisms including changes in the membrane lipid composition, changes in the phospholipid content [134], modifications of the cell surface charge, biofilm formation and various efflux pumps (for review see [13]. *Pseudomonas* spp. can degrade QACs under oxic conditions. A QAC tolerant and degrading *Pseudomonas fluorescence* strain was for example cultivated by enrichment cultivation with DADMAC-C10 as sole carbon source from activated sludge of a municipal WWTP [135]. *Pseudomonas* spp. were also most abundant in different BAC enrichment cultures with BACs as sole carbon and energy sources [32,130,136]. Sousa-Silva et al. [137] demonstrated in an exposure experiment with a *P. fluorescence* strain that the exposure to sublethal concentration with BACs contributes to the selection of strains with increased QAC tolerance. The authors also demonstrated that the adapted strains had an increased tolerance to the antibiotic ciprofloxacin. Here, 5% of the QAC tolerant *Pseudomonas* strains showed a ciprofloxacin resistance. Similarly, it was demonstrated for *P. aeruginosa* that the overexpression of efflux pumps leading to QAC resistance was responsible for a cross-resistance to ciprofloxacin. Harrison et al. [138] demonstrated in a mesocosm experiment with a freshwater bacterial community that the exposure to BACs (0.1 μg L^−1^ to 500 μg L^−1^) significantly increased the abundance of sulfamethoxazole and ciprofloxacin resistant bacteria. The cultivated bacteria were not further identified in that study. Kim et al. [125] demonstrated in a long-term exposure experiment of a river sediment bacterial community that exposure to BACs leads to the co-selection of ARBs. *P. aeruginosa* was among the ARBs cultivated after extended BAC exposure. Tandukar et al. [32] performed a long-term (4 years) enrichment experiment with contaminated river sediment using a mixture of BAC-C12 and -C14. They determined in enrichment cultures with BACs as sole C and energy source different *Pseudomonas* phylotypes as dominating taxa. None of the other taxa detected in their study as potential QAC degrading bacteria were cultivated in our study as BAC-C12 resistant bacteria. The authors showed in their study that BAC exposure decreased community diversity and increased AMR. We could also show that the diversity of bacteria cultured in the presence of BAC-C12 was reduced and that several of the cultivated strains contained additional antimicrobial resistance. We did however have no antimicrobial susceptibility test data on bacteria cultured in the absence of BAC-C12 and could not confirm if the MDR status increased in QAC tolerant strains.

While there are several studies available on the QAC tolerance of *Pseudomonas* spp. less is known for *Providencia* spp. It was so far only reported that *Providencia* strains were identified beside *Alcaligenes* strains as QAC tolerant bacteria cultivated from different WWTP effluents and freshwater habitats in Germany [139]. The genus *Providencia* belongs to the order *Enterobacteriales* and comprises facultative anaerobic mobile Gram-negative bacteria. It is next related to the genera *Proteus* and *Morganella* and contains a number of clinically relevant pathogenic species including *P. stuaetii*, *P. rettgeri,* and *P. alcalifaciens* [140]. Representatives of those species were cultivated from human faeces samples [140]. The here cultured *Providencia* strains showed highest 16S rRNA gene sequence identity to those species. For few strains class 1 integrons with *qacE* genes were detected, but QAC resistance mechanisms were so far not further described.

Finally, we would like to focus on some waste stream system specific results of cultivated bacteria. *Bacillus* species were cultivated only from livestock waste streams under non-selective cultivation conditions. The high abundance of cultivated *Bacillus* spp. agreed with previous studies of BGP input and output samples [141]. *Bacillus* spp. were however not cultivated in the presence of BAC-C12, which was different compared to other studies Forbes et al. [129] cultivated *Bacillus* species from a BAC exposed mesocosm experiment with kitchen drain biofilm communities and Gaze et al. [31] and Guo et al. [128] from agricultural soils on BAC containing agar media. It was shown by Brötzel and Cloetes [142] that *Bacillus cereus* and *Bacillus subtilis* strains can adapt to higher QAC concentrations by subsequent exposure to increasing QAC concentrations. Maybe the missing QAC tolerance of the *Bacillus* spp. strains present in livestock samples is due to the low concentrations of QACs present in those samples. *Brucella* and *Vibrio* species were cultivated as BAC-tolerant taxa exclusively from livestock waste streams. *Brucella* were also cultivated by Guo et al. [128] as QAC tolerant bacteria from contaminated water from shale gas production sites. QAC tolerance of *Brucella* (including now members of the genus *Ochrobactrum*) in waste stream systems was not yet studied. The *Brucella* strains cultivated here were all cultivated from farms 4 and 6. In contrast, *Vibrio* strains were cultivated from farms 1 and 2, the two frequently studied dairy farms. All cultivated *Vibrio* strains shared highest 16S rRNA gene sequence identity to the type strains of *V. cincinnatiensis.* The type strain of this species is an opportunistic human pathogen [143]. Several further strains of this genus were cultivated form domesticated animals including pigs and cattle in Germany and are associated to aborts [144]. The tolerance to QAC or biocidal compounds was not yet studied for this genus.

BAC-C12 tolerant *Aeromonas* strains were cultivated in our study only from human waste stream samples. All strains were next related to *A. hydrophila,* which is one of the common *Aeromonas* species that can cause animal and human infections as opportunistic pathogen [145]. *Aeromonas* spp. are known as typical freshwater bacteria frequently detected in various aquatic environments including freshwater, wastewater, and drinking water [145]. The absence of cultured *Aeromonas* spp. in sludge samples confirmed their free-living lifestyle in wastewater. Because *Aeromonas* are typical freshwater bacteria they have a strong potential to distribute resistances from human wastewaters into the receiving aquatic environment. *Aeromonas* spp. are well-known to colonize the gastrointestinal tract of different fish species and can thereby easily enter to the human food chain [145]. The differences in the QAC MIC values of *Aeromonas* strains cultivated in the absence and presence of BAC-C12 showed that there is a subfraction of *Aeromonas* strains present in human waste stream which had a reduced susceptibility to QACs. Few studies on environmental *A. hydrophila* strains with decreased BAC susceptibility are available. Confirming our data, Forbes et al. [129] cultivated *A. hydrophila* related strains in the presence of BACs and could show that the relative abundance of *Aeromonas* spp. increased during their long term QAC exposure mesocosm experiment. Patrauchan & Oriel [146] have shown that an *A. hydrophila* strain cultivated from a polluted soil was able to degrade BACs with an alkyl change length of 12, 14, and 18. An increased degradation ability was induced by the exposure to low BAC concentrations. Chacón et al. [147] performed a genome and proteome-based study of a less BAC susceptible *A. hydrophila* strain cultivated in the presence of BAC from a BAC-exposed activated sludge. The genome of the strain showed several mutations in genes linked to antimicrobial resistance and QAC tolerance including e.g. *qacC* and several efflux pump coding genes. Proteome studies showed that the main response of the strain to QAC exposure was an upregulation of several efflux pumps and downregulation of porines. While the strain of Chacón et al. [147] had no additional antibiotic resistances, our QAC tolerant strains partially exhibited a ciprofloxacin, cephalosporine, fosfomyin, chloramphenicol and TMP/SMX resistance. We cultivated one *A. hydrophila* related strain that carried a *qacE/qacEΔ1* gene together with *intI1* indicating that *Aeromonas* spp. seemed to be involved in the exchange of resistance determinants in human waste streams.

Beside *Aeromonas,* strains of few other taxa were exclusively cultivated as QAC tolerant bacteria from the human waste streams, namely *Citrobacter*, *Roultella*, *Novospirillum*, *Comamonas,* and *Klyvera* which include several genera that include species that harbour human pathogenic strains. The role of QAC-tolerance in those species is currently less explored.

## 5 Summary and conclusion

This study shows that QACs, QAC RGs, and viable QAC tolerant bacteria are released from both livestock and human waste streams into the environment via organic fertilisers (manure, BGP digestates, and biosolids) and treated wastewater (WWTP effluents). The release was in general higher for human compared to livestock waste streams as among the three common organic fertilizers used in agriculture, biosolids showed much higher loads of QACs, RGs, and fractions of QAC tolerant bacteria than manure and BGP digestates. Anaerobic digestion in BGPs showed a slight reduction, but no complete elimination, of these pollutants while in WWTPs, QACs and QAC tolerant bacteria seemed to accumulate in anaerobically digested biosolids. Anaerobic treatments therefore must be evaluated in a system-specific context. In human wastewater, a reduction of the absolute load of QACs, QAC RGs, and viable QAC tolerant bacteria was observed between the WWTP influent and effluent. In contrast, the relative abundance of QAC RGs and viable QAC tolerant bacteria increased, indicating a positive selection for QAC tolerance during the WWT process. This was also confirmed by higher QAC tolerance among abundant QAC tolerant bacteria cultivated from WWTPs compared to livestock systems, which was shown here for the first time

A clear indication for an increased MDR status of cultivated QAC tolerant bacteria was not abserved, but all QAC tolerant strains also carried antibiotic resistances. The presence of intI1 and *qacE/qacE1* genes in genetically different QAC tolerant strains indicates that there is at least a capacity for ARG uptake in genetic elements present in the QAC tolerant strains. More research is required to understand the interplay of QAC accumulation, adapted QAC tolerance and AMR spread in waste streams in more detail. The number of samples studied was too low to clearly demonstrate a pandemic effect; however, we were able to show that biosolids produced after the onset of the pandemic were enriched in both QACs and QAC-tolerant bacteria. This study furthermore showed that more prudent use of QACs and more efficient purification technologies will be required in the future to remove anthropogenic pollutants in human and livestock waste streams since purified wastewater and organic fertilizers will become increasingly important for sustainable agriculture due to the raising global temperatures, increasing water demand and the overuse of mineral fertilizers.

## CReDiT Contributions

Sophie Lennartz: Conceptualization, Data curation, Formal Analysis, Investigation, Methodology, Validation, Visualization, Writing – Original draft

Osas Elizabeth Aigbekaen: Data curation, Formal Analysis, Investigation, Methodology, Writing – Review and editing

Alexandra Jahraus: Data curation, Formal Analysis, Investigation

Jan Siemens: Resources, Funding acquisition, Writing – Review and editing

Ines Mulder: Conceptualization, Supervision, Funding acquisition, Validation, Writing – Review and Editing

Stefanie P. Glaeser: Conceptualization, Supervision, Data curation, Formal Analysis, Funding acquisition, Methodology, Validation, Visualization, Writing – Original draft

## Acknowledgements

This study was funded by the German Research Foundation (DFG) project numbers 458460392 and 431531292 and the Federal Ministry of Education and Research (BMBF)-funded JPI-EC-AMR JTC2017 Project ARMIS (Antimicrobial Resistance Manure Intervention Strategies, project number 01K11733). The procurement of the Liquid Chromatography Mass Spectrometer (HPLC-MS/MS) was funded jointly by the DFG (project number 428824233) and the Justus Liebig University Giessen. We thank Sanjana Balachandran for her great contributions to the microbiological analyses and Naomi Dietrich for assisting with QAC extractions.

