## Supplementary material for "Quaternary ammonium compounds (QACs), QAC resistance genes, and QAC tolerant bacteria in livestock and human waste streams": Supplementary Information_QAC_2025-12-16.pdf

*Shared first authors

### corresponding author

Contents

Text S1 Detailed description of QAC extraction procedures
Text S2 Method validation
Text S2.1 Recovery
Text S2.2 Matrix effect
Text S2.3 Limits of detection and quantification
Text S2.4 Linearity
Text S2.5 Precision
Table S1 Overview of studied livestock systems
Table S2 Overview of studied human WWTPs
Table S3 Average dry matter content of investigated samples
Table S4 QAC analytes and internal standards used in this study.
Table S5 QAC recovery rates for investigated matrices
Table S6 Recovery of internal standards in wastewater
Table S7 Matrix effect in sewage sludge and manure extracts
Table S8 Matrix effect in extracts of liquid wastewater
Table S9 Matrix effect in extracts of wastewater solids
Table S10 Instrumental intraday precision
Table S11 Retention times and MRM parameters for QACs and internal standards.
Table S12 HPLC and MS settings
Table S13 Primer systems used for quantitative PCRs (qPCRs) and generation of qPCR standards
Table S14 Standards and qPCR conditions
Table S15 Overview of QAC, qPCR, and CFU data obtained for studied livestock and waste stream samples
-> see Excel table as separate document

Table S16 Wilcoxon rank sum test between cattle vs. pig manure and BGP digestates

Table S17 QAC distribution between the liquid and particulate phases of influent, activated sludge and effluent

Tables S18-S25 Statistical results

-> see Excel table as separate document

Supplementary Figures

Fig. S1 Proportion of ATMAC-C16, BAC-C12 and DADMAC-C10 in the total QAC concentrations (excluding DADMAC-C18 and CLQ)
Fig. S2 Phase distribution of QACs in influent and activated sludge
Fig. S3 QAC mass balances in the rural and urban WWTP
Fig. S4 Spearman correlations between dry weight concentrations of QACs, the fraction of BAC-C12 tolerant bacteria, and relative gene abundances among all analysed (semi-)solid samples (manure, biogas plant digestate, activated sludge, dewatered sludge and anaerobically digested sludge; n=30).
Fig. S5 Spearman correlations between the fraction of BAC-C12 tolerant bacteria and relative gene abundances among all analysed samples (manure, biogas plant digestate, activated sludge, dewatered sludge, anaerobically digested sludge, WWTP influent, WWTP effluent; n=37).
Fig. S6 Comparative analysis of BAC-C12 and DADMAC-C10 MIC distribution patterns among strains of the three most abundant genera cultivated in the absence and presence of BAC-C12 (*Aeromonas, Escherichia/Shigella,* and *Alcaligenes*).
Fig. S7 Comparative analysis of BAC-C12 and DADMAC-C10 MIC distribution patterns for *Providencia* and *Pseudomonas* strains separated based on individual sampling types of both waste stream systems.

Table S3 Overview of studied livestock systems. Modified based on Pulami et al. [1].

| Farms and respective BPGs | Farm type | Manure source | Co-substrates used in BGPs | BGP process conditions | BGP operating temperature | Sampling points pre/post pandemic |
|---|---|---|---|---|---|---|
| Farm1/ BGP1 | Dairy farming | Dairy cattle | Corn, grain/cereal, grass | Mesophilic | ~40°C | 2/2 |
| Farm2/ BGP2 | Dairy farming | Dairy cattle | Corn, grain/cereal | Thermophilic | ~50°C | 2/2 |
| Farm4/ BGP4 | Meat Production | Pigs | Corn, grain/cereal | Mesophilic | ~40°C | 1/0 |
| Farm5/ BGP5 | Beef production | Pigs, beef, chicken | Maize, green rye, sugarbeet, cereal meal, pig/cattle slurry | Mesophilic | ~40°C | 1/0 |
| Farm6/ BGP6 | Dairy farming | Dairy cattle | Maize, green rye, sugarbeet, cereal meal, pig slurry | Mesophilic | ~40°C | 1/0 |

Table S4 Overview of studied human WWTPs

| WWTPs | Catchment area | Treatment type | Inhabitant equivalents | Capacity | Sludge production (dry weight) | Sampling dates pre/post pandemic |
|---|---|---|---|---|---|---|
| WWTP1 | Rural (no medical facilities) | Biological treatment, mixed tank with altered aerobic and anaerobic phases; no anaerobic sludge treatment | 13,000 | 39 L $s^{-1}$ (average 2019-2021) | 174 t $a^{-1}$ (average 2019-2021) | 2/1 |
| WWTP6 | Urban (17 human and veterinarian hospitals/ clinics) | Biological treatment, separated aerobic and anaerobic treatment areas; anaerobic sludge treatment | 280,000 | 710 L $s^{-1}$ (2021 average) | 3,350 t $a^{-1}$ (2021) | 0/1* |

* QAC concentrations in 2016 are available from Heyde et al. [2]

Table S3 Average dry matter content of investigated samples

| Dry weight (mass %) | Livestock farms | Rural WWTP (WWTP1) | Urban WWTP (WWTP6) |
|---|---|---|---|
| Raw manure (M) | 8.93 | - | - |
| BGP digestates (BGP-D) | 8.4 | - | - |
| Influent (I) | - | 0.025 | 0.022 |
| Activated sludge (AS) | - | 0.66 | 0.58 |
| Dewatered sludge (S) | - | 26.51 | - |
| Anaerobically digested sludge (ADS) | | - | 26.33 |
| Effluent (E) | - | ~0.01 | <0.01 |

Table S4 QAC analytes and internal standards used in this study. The analytes had a chemical purity between 95-99% and the internal standards between 92-99%.

| Compound name | Abbreviation | CAS number | Supplier |
|---|---|---|---|
| Alkyltrimethylammonium compounds | ATMACs | | |
| Octyltrimethyl ammonium bromide | ATMAC-C8 | 2083-68-3 | TCI* |
| Decyltrimethyl ammonium chloride | ATMAC-C10 | 10108-87-9 | TCI* |
| Dodecyltrimethyl ammonium chloride | ATMAC-C12 | 112-00-5 | TCI* |
| Tetradecyltrimethyl ammonium chloride | ATMAC-C14 | 4574-04-3 | TCI* |
| Hexadecyltrimethyl ammonium chloride | ATMAC-C16 | 112-02-7 | TCI* |
| Benzylaklylammonium compounds | | | |
| Benzyldimethyloctylammonium chloride | BAC-C8 | 959-55-7 | Sigma Aldrich‡ |
| Benzyldimethyldecylammonium chloride | BAC-C10 | 965-32-2 | Sigma Aldrich‡ |
| Benzyldimethyldodecylammonium chloride | BAC-C12 | 139-07-1 | TCI* |
| Benzyldimethyltetradecylammonium chloride | BAC-C14 | 139-08-2 | TCI* |
| Benzyldimethylhexadecylammonium chloride | BAC-C16 | 122-18-9 | TCI* |
| Benzyldimethyloctadecylammonium chloride | BAC-C18 | 122-19-0 | TCI* |
| Dialkyldimethylammonium compounds | | | |
| Dioctyldimethylammonium bromide | DADMAC-C8 | 3026-69-5 | TCI* |
| Dioctyldidecylammonium chloride | DADMAC-C10 | 7173-51-5 | TCI* |
| Dioctyldidodecylammonium bromide | DADMAC-C12 | 3401-74-9 | TCI* |
| Dioctylditetradecylammonium bromide | DADMAC-C14 | 68105-02-2 | TCI* |
| Dioctyldihexadecylammonium chloride | DADMAC-C16 | 70755-47-4 | TCI* |
| Dioctyldioctadecylammonium chloride | DADMAC-C18 | 107-64-2 | Sigma Aldrich‡ |
| Other QACs | | | |
| Benzethonium chloride | BEC | 121-54-0 | Sigma Aldrich‡ |
| Chlormequat chloride | CLQ | 999-81-5 | Sigma Aldrich‡ |
| Internal standards | | | |
| D7- Benzyldimethyloctylammonium chloride | BAC-C8-$D_7$ | -§ | HPC Standards† |
| D7- Benzyldimethyldodecylammonium chloride | BAC-C12- $D_7$ | -§ | HPC Standards† |
| D7- Benzyldimethyloctylammonium chloride | BAC-C14- $D_7$ | 1219178-72-9 | HPC Standards† |
| D7- Benzyldimethyloctadecylammonium chloride | BAC-C18- $D_7$ | -§ | HPC Standards† |
| D6-Dioctyldimethylammonium iodide | DADMAC-C8-$D_6$ | -§ | HPC Standards† |
| D6-Dioctyldimethylammonium chloride | DADMAC-C10-$D_6$ | -§ | HPC Standards† |
| D6-Didodecyldimethylammonium iodide | DADMAC-C12-$D_6$ | -§ | HPC Standards† |
| D6-Ditetradecyldimethylammonium iodide | DADMAC-C14-$D_6$ | -§ | HPC Standards† |

**Eschborn, Germany*

*‡ Steinheim, Germany*

*† Cunnersdorf, Gemany*

§ no separate CAS number from unlabeled compound

Text S1: Detailed description of QAC extraction procedures

S1.1 Sewage sludge, manure, and BGP digestates

Before extraction, the samples were lyophilized and carefully homogenized with porcelaine mortar and pestle. Due to coarse straw and wood residues in the BGP input samples, preliminary tests yielded very low recoveries and poor reproducibility. To improve extractability and sample homogeneity, all BGP samples were consequently milled to fine powder using a planetary ball mill equipped with agate grinding cups ("Pulverisette 5", Fritsch GmbH, Idar-Oberstein; settings 400 rpm x 2,5 min, 4 cycles with 12 balls). For QAC extraction, 10 mL of acidified acetonitrile (0.1 vol% HCl) was added to a 1.5-2 g sample aliquot and three extraction cycles of 10 min shaking at 250 rpm on a horizontal shaker, 10 min ultrasonication at room temperature, and 10 min centrifugation at 2000 rpm (Rotana 460R, Hettich, Tuttlingen, Germany) were applied. The supernatant was collected after each cycle, pooled, and concentrated to < 1 mL in a Syncore Polyvap evaporator unit (Büchi, Flawil, Switzerland). Afterwards, the extract was diluted with 9 mL ultrapure water and loaded onto 500 cc chromabond CN solid-phase extraction (SPE) cartridges (Machery Nagel, Düren, Germany) pre-conditioned with 6 mL acetonitrile and 12 mL milli-Q water. The cartridges were washed with milli-Q water and 1M HCl before eluting QACs with 2 mL acidifed methanol (4:1 methanol/1M HCl) into 15 mL centrifuge tubes (Th. Geyer GmbH & Co. KG, Renningen, Germany) as described by Heyde et al. (2020). The eluted sampled were dried under nitrogen at 40 °C, reconstituted in 1 mL acetonitrile, and centrifuged at 4000 rpm (Rotanta 460R) for 30 min to precipitate remaining particles. The particle-free supernatant was transferred into HPLC vials (neoLab Migge, Heidelberg, Germany) and stored at -20°C until analysis.

S1.2 WWTP influent, activated sludge, and effluent

Influent, activated sludge and effluent were separated into operationally defined "solid" and "liquid" phases. The activated sludge samples, which had a high particle content (5.4-7.4 g per L), were centrifuged in the sampling vessels immediately after return from the WWTPs. The liquid supernatants were separated from the solid pellets and both were frozen until further analysis. The pellets were freeze-dried, weighed into amber glass vials, and extracted as described above (S1.1). Weighing of the original wastewater sample and lyophilized pellet after centrifugation allowed for back-calculation of the approximate dry mass fraction, neglecting the contribution of non-settleable particles.

For analysis of the liquid phase of the influent, activated sludge and effluent, a 150-200 mL aliquot was thawed, homogenized by swirling, divided between 3-4 50 mL centrifuge tubes, and centrifuged for 30 min at 4,160 rpm (Rotana 460R, Hettich, Tuttlingen, Germany). This step was necessary for all matrices to avoid blockage of the SPE cartridges. The liquid supernatants were pooled back into one glass beaker, fortified with 200 ng of BAC-C14-$D_7$ and DADMAC-C14-$D_6$ as surrogate standards, and pH-adjusted with 30 µL $NH_4OH$ per 200 mL (~pH10) for improved SPE recovery as suggested by Östman et al. (2017). Oasis HLB 150 cc cartridges were conditioned with 5 mL methanol and 5 mL milli-Q before loading the samples over 70 mL PTFE reservoirs (Merck, Darmstadt, Germany) onto the SPE cartridges. The cartridges were washed with 3mL milli-Q water and QACs were eluted with 5 mL acidified methanol (0.1 vol% formic acid) and 5 mL ethyl acetate. The eluates were dried under nitrogen, resuspended in acetonitrile, and transferred to HPLC vials. The remaining particulate phase of the influent and effluent was freeze-dried and then processed directly in the falcon tubes following the shaking-ultrasonic procedure described in S1.1. Due to the low particle yield, no SPE clean-up was necessary. Instead, the concentrated extracts were transferred to 2 mL safelock tubes, reconstituted in 250 µL ACN and then centrifuged for 15 min at 17,000 g to remove residual particles. The final 250 µL extracts were transferred to 450 µL glass inserts in HPLC vials and frozen until analysis.

Text S2 Method validation

S2.1 Recovery

S2.1.1 Biosolids extraction method

To assess QAC recovery rates from biosolids, roughly 10 g aliquots of manure, BGP digestates or dewatered sludge were covered with pentane in a 250 mL Schott flask and spiked with the QAC standard mix. The BGP samples, which had relatively low QAC background levels, were spiked with 50–100 ng g d.w.$^{-1}$. The spiking concentrations for sewage sludge were 500 ng g$^{-1}$ for the rural WWTP and 1000 ng g$^{-1}$ for the urban WWTP, which corresponds to roughly 50% of the measured concentration of the most abundant QAC homologue. After shaking for 24 hours on an overhead shaker, the flasks were opened, and the pentane was evaporated under repeated stirring. Triplicates of 2g of dried spiked samples were then extracted along with unspiked samples a few days after the spiking to mimic aging. For each manure, BGP digestates and dewatered sewage sludge, two representative samples were tested. A method blank spiked with 100 ng of each QAAC was processed alongside routine extractions to determine recovery without matrix interferences. Recovery rates were calculated from the background-subtracted concentration in the spiked samples relative to the nominal concentration as:

$$\text{Recovery (\%)} = (\text{conc}_{\text{spiked sample}} - \text{conc}_{\text{sample}}) / \text{conc}_{\text{nominal}} \times 100$$

Recovery rates of different QACs from the spiked blank ranged from 5-89% (median 43%) when including the SPE cleanup (Table S5), and from 40-70% without SPE (data not shown), which is similar to results for the original method (Heyde et al., 2020). Recoveries were highest for the short-and single-chained QACs but very low (<10%) and highly variable for CLQ and DADMAC-C18, independent of the spike level and analyzed matrix. The hydrophilic chlormequat was probably removed with the aqueous SPE washing solution while the rather hydrophobic DADMAC-C18 may have only been weakly retained on the cyanopropyl SPE phase or not efficiently eluted with methanol. The applied extraction method was therefore not suitable for CLQ and DADMAC-C18 and reported concentrations in the supplementary data file are only semi-quantitative.

Matrix-dependent differences in recovery rates were observed between BGP and WWTP biosolids (Table S5). For spiked BGP solids, recoveries ranged from 5-39% (median 26%, excluding CLQ and DADMAC-C18) and for spiked sewage sludge from 11- >100% (median 57%). Particularly BACs with alkyl chains ≥C16 and DADMACs with chain length ≥C12 showed very low recoveries (<15%) from spiked manure and BGP digestates. Non-detects for CLQ, BACs C16-C18 and DADMACs C12 to C-18 in livestock samples may therefore be due to underestimation of actual concentrations. Recoveries from sewage sludge were overall higher, but in some cases more uncertain due to higher background concentration. For example, recoveries >100% were calculated for ATMAC-C16 and DADMAC-C10 despite subtracting background levels; however, these are not considered accurate as they did not occur in blank spikes. Differences in the recovery rates between BGP and WWTP biosolids may also be attributable to the 10-fold lower spiking level of the farm solids, causing a higher proportion of QACs to be lost on surfaces during the sample cleanup, or different degrees of matrix interference due to the different sample composition.

S2.1.2 Wastewater

Recovery from the liquid wastewater phase could not be evaluated from spiked samples due to the limited available sample volumes. Therefore, recovery was assessed using triplicates of 150 mL matrix-free QAC solutions in ultrapure water (spiked to a final concentration of 5 and 500 ng L$^{-1}$ for each homologue). Residues in both the liquid phase and losses due to adsorption to the extraction tubes were calculated. The actual recovery rates for the wastewater particles were assessed by performing recovery trials with 0.5 g of riverine suspended particulate matter spiked with 200 ng g$^{-1}$ of QACs.

Due to potential impact of the wastewater matrix on recovery, two mid-chained internal standards (IS), BAC-C14-$D_7$ and DADMAC-C14-$D_6$, were added as extraction standards into the liquid WWTP samples prior to SPE. Although not representative for all QAC homologues, this allowed to estimate differences in recovery between influent, activated sludge and effluent samples (Table S9). Recovery of the IS was calculated as:

$$\text{Recovery (\%)} = (\text{area}_{\text{spiked method blank}} - \text{area}_{\text{method blank}}) / (\text{area}_{\text{spiked solvent blank}}) \times 100$$

QAC recoveries for the liquid wastewater phase ranged from 6-96% (median 57% and 62% depending on the spiking concentration), excluding unrealistic recoveries < 0 for DADMAC-C18 and BAC-C18 at 500 ng $L^{-1}$ and >> 100% for BAC-C12 and BEC at 5 ng $L^{-1}$. Recoveries decreased with increasing chain length of QACs, especially for homologues ≥ C16 and DADMACs (Table S5). No more than 6% of the spiked concentrations was recovered from the extraction tubes (data not shown), indicating that the calculated solid-phase concentrations originate primarily from particle-attached QACs and only to a minor extent from sorption of liquid-phase QACs to the labware. Extraction tubes contained higher amounts of QAC homologues with longer or two alkyl chains, suggesting that surface adsorption increased with stronger hydrophobicity of QACs. Between 14-79% (median 56%) of spiked QACs were recovered from river particles used as a proxy for wastewater particles (Table S5).

Table S5 QAC recovery rates for investigated matrices. * indicates unrealistic values due to high background

| Extraction method | Manure & sludges | | | | | Wastewater | | |
|---|---|---|---|---|---|---|---|---|
| Matrix | Blank | Manure | BGP-D | Dewatered sludge WWTP-1 | Anaerobically digested sludge WWTP-6 | Liquid | | Solid |
| Spike level | 100 ng | 50-100 ng $g^{-1}$ | 50-100 ng $g^{-1}$ | 0.5 µg $g^{-1}$ | 1 µg $g^{-1}$ | 5 ng $L^{-1}$ | 500 ng $L^{-1}$ | 200 ng $g^{-1}$ |
| Replicates | 3 | 9 | 6 | 3 | 3 | 3 | 3 | 4 |
| ATMAC-C8 | 80 ± 7 | 35 ± 5 | 32 ± 6 | 86 ± 13 | 40 ± 3 | 85 | 81 | 50 |
| ATMAC-C10 | 82 ± 10 | 33 ± 7 | 30 ± 7 | 73 ± 3 | 49 ± 2 | 97 | 82 | 53 |
| ATMAC-C12 | 78 ± 13 | 27 ± 9 | 27 ± 8 | 72 ± 2 | 56 ± 2 | 85 | 76 | 53 |
| ATMAC-C14 | 73 ± 11 | 28 ± 10 | 27 ± 8 | 62 ± 3 | 52 ± 2 | 57 | 55 | 56 |
| ATMAC-C16 | 50 ± 13 | 28 ± 14 | 27 ± 11 | 59 ± 7 | 128 ± 13* | 28 | 29 | 58 |
| BAC-C8 | 89 ± 12 | 42 ± 6 | 39 ± 7 | 55 ± 4 | 53 ± 1 | 94 | 89 | 56 |
| BAC-C10 | 75 ± 15 | 25 ± 10 | 26 ± 8 | 49 ± 3 | 65 ± 2 | 86 | 70 | 59 |
| BAC-C12 | 68 ± 19 | 24 ± 11 | 26 ± 10 | 57 ± 6 | 35 ± 161* | 74 | 245* | 58 |
| BAC-C14 | 40 ± 8 | 19 ± 10 | 19 ± 8 | 49 ± 8 | 96 ± 48* | 41 | 74 | 48 |
| BAC-C16 | 16 ± 6 | 12 ± 6 | 12 ± 5 | 82 ± 1 | 53 ± 13 | 24 | 18 | 48 |
| BAC-C18 | 27 ± 14 | 9 ± 15 | 9 ± 12 | 86 ± 5 | 38 ± 8 | 15 | -230* | 48 |
| DADMAC-C8 | 81 ± 10 | 38 ± 15 | 38 ± 13 | 69 ± 3 | 11 ± 11 | 94 | 74 | 57 |
| DADMAC-C10 | 24 ± 6 | 19 ± 12 | 19 ± 10 | 53 ± 39 | 968 ± 299* | 38 | 70 | 33 |
| DADMAC-C12 | 27 ± 15 | 15 ± 10 | 14 ± 8 | 65 ± 1 | 7 ± 1 | 16 | 16 | 60 |
| DADMAC-C14 | 37 ± 9 | 13 ± 9 | 12 ± 7 | 68 ± 2 | 38 ± 1 | 27 | 10 | 79 |
| DADMAC-C16 | 27 ± 10 | 5 ± 4 | 5 ± 3 | 11 ± 2 | 12 ± 3 | 32 | 6 | 69 |
| DADMAC-C18 | 9 ± 6 | 1 ± 1 | 1 ± 1 | -2.4 ± 2 | 4 ± 5 | 21 | -7* | 860* |
| BEC | 56 ± 7 | 25 ± 11 | 26 ± 9 | 55 ± 5 | 59 ± 3 | 57 | 2939* | 53 |
| CLQ | 5 ± 2 | 1 ± 1 | 1 ± 1 | 1 ± 0.2 | 0.2 ± 0.1 | 21 | 16 | 14 |

Table S6 Recovery of internal standards from wastewater

| IS recovery | | n | BAC-C14-$D_7$ | | | DADMAC-C14-$D_6$ | | |
|---|---|---|---|---|---|---|---|---|
| WWTP-2 | AS | 3 | 24 | ± | 5 | 15 | ± | 3 |
| | E | 3 | 33 | ± | 4 | 63 | ± | 9 |
| | I | 2 | 13 | ± | 2 | 37 | ± | 9 |
| WWTP-3 | AS | 2 | 19 | ± | 0 | 37 | ± | 0 |
| | E | 3 | 34 | ± | 4 | 65 | ± | 5 |
| | I | 3 | 7 | ± | 0 | 47 | ± | 4 |

S2.2 Matrix effect

Matrix effects on the analytes ($ME_{analyte}$) were determined by post-extraction spiking with 200 ng of each QAC for sewage sludge and 100 ng for the other matrices. These concentrations were selected to approximate realistic sample concentrations that still fell in the linear calibration range without sample dilution. The $ME_{analyte}$ was calculated as percentage of ionization suppression or enhancement:

$$ME_{analyte}\ (\%) = (area_{spiked\ sample} - area_{sample}) / (area_{spiked\ blank} - area_{blank}) \times 100 - 100.$$

Thus, values below or above zero indicate signal suppression or enhancement, respectively. The matrix effect on the internal standards ($ME_{IS}$) was also routinely determined based on the IS area at in each sample relative to the average IS area in the calibrants:

$$ME_{IS}\ (\%) = (area_{\ IS,\ sample}) / (mean\ area_{\ IS,\ calibrant}) \times 100 - 100.$$

The blank subtraction was not necessary since the internal standards were not present in unspiked samples or pure solvent blanks.

Strong analyte signal suppression was observed in extracts of dewatered sewage sludge (-29 to -83%) and manure (-5 to -67%; Table S7). In liquid wastewater extracts, most analytes showed signal suppression as well while the signal of DADMACs and CLQ was occasionally enhanced (Table S8). In extracts of the particulate phase of the effluent, the signal of all analytes and IS was enhanced (+12 to +48% and +12 to +27%, respectively), while extracts of the influent particles showed both signal suppression and enhancement (Table S9).

The internal standards mostly showed similar behavior as the analytes and thus compensated for the ME to a large extent (Tables S7-S9). However, they performed less well for the longer chained DADMACs, probably due to the larger difference between analyte and IS retention time for these compounds. In cases where the analyte concentrations were outside the linear calibration range, the calculated ME may not be accurate (e.g. >100% signal suppression was calculated for BAC-C12 and DADMAC-C10 in sludge; Table S7).

Table S7 Matrix effect on analytes and internal standards (IS, 20 ng $mL^{-1}$) in sludge and manure extracts. * no reliable quantification due to high background analyte levels.

| STD | IS | Sludge | | | Manure | | |
|---|---|---|---|---|---|---|---|
| | | STD | IS | diff | STD | IS | diff |
| ATMAC-C8 | BAC-C8-D7 | -44 | -41 | -3 | -59 | -52 | -8 |
| ATMAC-C10 | BAC-C8-D7 | -54 | -41 | -13 | -62 | -52 | -10 |
| ATMAC-C12 | DADMAC-C8-D6 | -54 | -60 | 7 | -61 | -55 | -6 |
| ATMAC-C14 | DADMAC-C8-D6 | -55 | -60 | 5 | -59 | -55 | -3 |
| ATMAC-C16 | DADMAC-C10-D6 | -71 | -79 | 8 | -49 | -63 | 14 |
| BAC-C8 | BAC-C8-D7 | -37 | -41 | 4 | -52 | -52 | 0 |
| BAC-C10 | BAC-C12-D7 | -43 | -47 | 5 | -48 | -37 | -12 |
| BAC-C12 | BAC-C12-D7 | -110* | -47 | -63 | -67 | -37 | -30 |
| BAC-C14 | BAC-C12-D7 | -83 | -50 | -33 | -64 | -47 | -17 |
| BAC-C16 | BAC-C18-D7 | -75 | -31 | -45 | -63 | -26 | -37 |
| BAC-C18 | BAC-C18-D7 | -70 | -31 | -40 | -47 | -26 | -21 |
| DADMAC-C8 | DADMAC-C8-D6 | -81 | -60 | -21 | -46 | -55 | 9 |
| DADMAC-C10 | DADMAC-C10-D6 | -149* | -79 | -70 | -78 | -63 | -15 |
| DADMAC-C12 | DADMAC-C12-D6 | -51 | -18 | -33 | -43 | -18 | -25 |
| DADMAC-C14 | DADMAC-C12-D6 | -29 | 3 | -32 | -17 | 3 | -20 |
| DADMAC-C16 | DADMAC-C12-D6 | -48 | 3 | -50 | -5 | 3 | -8 |
| DADMAC-C18 | DADMAC-C12-D6 | -83 | 3 | -86 | -26 | 3 | -29 |
| BEC | BAC-C12-D7 | -57 | -50 | -7 | -55 | -45 | -10 |
| CLQ | BAC-C8-D7 | -52 | -41 | -11 | -39 | -52 | 12 |

Table S8 Matrix effect on the analytes (STD) and internal standards (IS, 20 ng $mL^{-1}$) in extracts of liquid wastewater

| Matrix effect (%) | | WWTP-I (liquid) | | | WWTP-AS (liquid) | | | WWTP-E (liquid) | | |
|---|---|---|---|---|---|---|---|---|---|---|
| STD | IS | STD | IS | diff | STD | IS | diff | STD | IS | diff |
| ATMAC-C8 | BAC-C8-$D_7$ | -67 | -72 | 5 | -19 | -53 | 34 | -37 | -33 | -4 |
| ATMAC-C10 | BAC-C8-$D_7$ | -70 | -72 | 2 | -41 | -53 | 12 | -46 | -33 | -13 |
| ATMAC-C12 | DADMAC-C8-$D_6$ | -83 | -86 | 4 | -42 | -64 | 22 | -47 | -59 | 11 |
| ATMAC-C14 | DADMAC-C8-$D_6$ | -85 | -86 | 1 | -31 | -64 | 33 | -51 | -59 | 8 |
| ATMAC-C16 | DADMAC-C10-$D_6$ | -81 | -89 | 7 | -25 | -51 | 26 | -45 | -48 | 2 |
| BAC-C8 | BAC-C8-$D_7$ | -75 | -72 | -4 | -19 | -53 | 34 | -25 | -33 | 8 |
| BAC-C10 | BAC-C12-$D_7$ | -76 | -83 | 7 | -19 | -58 | 39 | -33 | -42 | 9 |
| BAC-C12 | BAC-C12-$D_7$ | 23 | -83 | 106 | -24 | -58 | 34 | -49 | -42 | -6 |
| BAC-C14 | BAC-C12-$D_7$ | -73 | -83 | 10 | -34 | -58 | 24 | -52 | -42 | -10 |
| BAC-C16 | BAC-C18-$D_7$ | -84 | -73 | -11 | -14 | -32 | 18 | -30 | -20 | -10 |
| BAC-C18 | BAC-C18-$D_7$ | -65 | -73 | 8 | 12 | -32 | 43 | 0 | -20 | 20 |
| DADMAC-C8 | DADMAC-C8-$D_6$ | -13 | -86 | 74 | -20 | -64 | 45 | -34 | -59 | 24 |
| DADMAC-C10 | DADMAC-C10-$D_6$ | -51 | -89 | 38 | -41 | -51 | 10 | -57 | -48 | -9 |
| DADMAC-C12 | DADMAC-C12-$D_6$ | -54 | -59 | 5 | 13 | -14 | 27 | 1 | -5 | 6 |
| DADMAC-C14 | DADMAC-C12-$D_6$ | 19 | -59 | 78 | 37 | -14 | 51 | 20 | -5 | 25 |
| DADMAC-C16 | DADMAC-C12-$D_6$ | 108 | -59 | 167 | 69 | -14 | 83 | 48 | -5 | 54 |
| DADMAC-C18 | DADMAC-C12-$D_6$ | 102 | -59 | 161 | 75 | -14 | 89 | 58 | -5 | 64 |
| BEC | BAC-C12-$D_7$ | -86 | -83 | -3 | -20 | -58 | 38 | -43 | -42 | 0 |
| CLQ | BAC-C8-$D_7$ | -49 | -72 | 23 | 15 | -53 | 69 | -20 | -33 | 13 |

Table S9 Matrix effect on the analytes (STD) and internal standards (IS, 20 ng $mL^{-1}$) in extracts of wastewater solids

| Matrix effect (%) | | WWTP-I solids | | | WWTP-E solids | | |
|---|---|---|---|---|---|---|---|
| STD | IS | STD | IS | diff | STD | IS | diff |
| ATMAC-C8 | BAC-C8-$D_7$ | 0 | -7 | 7 | 30 | 12 | 18 |
| ATMAC-C10 | BAC-C8-$D_7$ | 0 | -7 | 6 | 25 | 12 | 13 |
| ATMAC-C12 | DADMAC-C8-$D_6$ | 1 | -23 | 23 | 25 | 17 | 8 |
| ATMAC-C14 | DADMAC-C8-$D_6$ | -16 | -23 | 7 | 26 | 17 | 9 |
| ATMAC-C16 | DADMAC-C10-$D_6$ | 33 | -31 | 64 | 26 | 13 | 13 |
| BAC-C8 | BAC-C8-$D_7$ | 6 | -7 | 12 | 26 | 12 | 13 |
| BAC-C10 | BAC-C12-$D_7$ | 3 | -16 | 19 | 23 | 20 | 3 |
| BAC-C12 | BAC-C12-$D_7$ | -78 | -16 | -62 | 12 | 20 | -8 |
| BAC-C14 | BAC-C12-$D_7$ | -70 | -16 | -55 | 18 | 20 | -2 |
| BAC-C16 | BAC-C18-$D_7$ | -11 | -1 | -9 | 24 | 21 | 3 |
| BAC-C18 | BAC-C18-$D_7$ | -5 | -1 | -3 | 22 | 21 | 1 |
| DADMAC-C8 | DADMAC-C8-$D_6$ | 31 | -23 | 53 | 24 | 17 | 8 |
| DADMAC-C10 | DADMAC-C10-$D_6$ | 154 | -31 | 185 | 16 | 13 | 3 |
| DADMAC-C12 | DADMAC-C12-$D_6$ | -14 | 7 | -22 | 22 | 27 | -5 |
| DADMAC-C14 | DADMAC-C12-$D_6$ | -2 | 7 | -10 | 28 | 27 | 1 |
| DADMAC-C16 | DADMAC-C12-$D_6$ | 39 | 7 | 31 | 43 | 27 | 16 |
| DADMAC-C18 | DADMAC-C12-$D_6$ | 84 | 7 | 76 | 48 | 27 | 21 |
| BEC | BAC-C12-$D_7$ | -12 | -16 | 4 | 23 | 20 | 3 |
| CLQ | BAC-C8-$D_7$ | -18 | -7 | -11 | 30 | 12 | 17 |

S2.3 Limits of detection and quantification

Instrumental limits of detection (IDL) and quantification (IQL) were determined from the analyte concentrations in solvent blanks as:

$$IDL = mean_{blank} + 3 * standard\ deviation_{blank}$$

$$IQL = mean_{blank} + 5 * standard\ deviation_{blank}$$

Blanks measured immediately following highly concentrated samples were omitted from the IDL and IQL calculation due to low-level carry-over. For each extraction procedure, the method limits of detection (MDL) and quantitation (MDL) were calculated analogously from the procedural blanks across all extraction days.

IDLs for all QACs ranged from 1-8 ng $mL^{-1}$ (median 2 ng $mL^{-1}$). MDLs for processed sludge, manure and BGP digestates fell between 1-22 ng $g^{-1}$ (median 2 ng $g^{-1}$) and were highest for ATMAC-C12, ATMAC-C8 and DADMAC-C10. Separate values apply for the activated sludge, which was first separated from the liquid phase by centrifugation. MDLs ranged from 0.2-4 ng $g^{-1}$ for most QACs while ATMAC-C16, BAC-C12 and DADMAC-C10 had notably higher MDLs of 98, 32 and 116 ng $g^{-1}$, respectively. Excessive blank contamination was probably caused by QAC residues on lab equipment and could be prevented during consequent extractions by more thorough cleaning of reusable glassware and SPE adapters. However, due to the low amount of sampling material the analysis of the activated sludge samples could not be repeated. MDLs for the liquid wastewater phase ranged from 2-293 ng $L^{-1}$ (median 8 ng $L^{-1}$) and for the solid wastewater phase between 1-61 ng $L^{-1}$ (median 7 ng $L^{-1}$). BEC, DADMAC-C10, DADMAC-C18 and BAC-C12 showed the highest levels of blank contamination in both phases. When comparing QAC concentrations, it is important to keep in mind that different MDLs apply for each matrix and QAC homologues.

#### S2.4 Linearity

Calibration curves of the area/ IS area ratio were visually assessed for linearity. If linearity was not given over the entire range from 0.5–200 ng $mL^{-1}$ for a particular analyte, the calibration curve was split to include a minimum of 5 levels in both the lower and higher concentration range [3]. The accuracy acceptance for calibrants was 80-120% and 70-130% for the lower limit of quantification. Overall, the fit of the linear model was very good for all QACs with coefficients of determination ($R^2$) > 0.98 for DADMAC-C18 and ≥ 0.99 for all other QACs.

#### S2.5 Precision

Measurement repeatability (intraday precision) was assessed for the analyte and IS signal in nine repeat injections of a low (1 ng $mL^{-1}$), medium (10 ng $mL^{-1}$) and high (100 ng $mL^{-1}$) concentration of the standard solution and in two matrix-rich samples including one manure sample and one dewatered sludge sample (n=8 injections).

Intraday precision of the analyte to IS area ratio of the different QACs in standard solution was concentration-dependent and ranged from 3-9% at 1 ng $mL^{-1}$, 1-7% at 10 ng $mL^{-1}$ and 1-4% at 100 ng $mL^{-1}$ (Table S10). It was slightly better than precision of only the analyte signal alone (3-12% RSD across all spiking levels; data not shown). Intraday precision was fairly good for all homologues in the spiked manure and dewatered sludge samples with RSDs between 2-9% (median 4.7%) for the analyte signal and 2-6% (median 3.1%) for the area ratio. CLQ was an exception with RSDs between 18-21% for the analyte signal and 7-8% for the area ratio, which may be due to its very early elution along with hydrophilic impurities and its low sample concentration. These results show that use of the IS was helpful in increasing precision both in pure standards and matrix-rich samples.

Table S10 Instrumental intraday precision (n=9 for standards, n=8 for spiked matrix)

| QAC | 1 ng $mL^{-1}$ | 10 ng $mL^{-1}$ | 100 ng $mL^{-1}$ | Manure (100 ng $mL^{-1}$) | WWTP-S (200 ng $mL^{-1}$) |
|---|---|---|---|---|---|
| ATMAC-C8 | 3.8 | 3.0 | 2.8 | 2.6 | 3.0 |
| ATMAC-C10 | 3.1 | 1.5 | 2.9 | 3.3 | 3.8 |
| ATMAC-C12 | 4.4 | 4.2 | 2.1 | 4.1 | 3.2 |
| ATMAC-C14 | 4.5 | 2.9 | 2.6 | 2.8 | 4.6 |
| ATMAC-C16 | 4.0 | 3.8 | 3.7 | 3.2 | 3.5 |
| BAC-C8 | 3.4 | 2.7 | 3.3 | 2.7 | 3.1 |
| BAC-C10 | 3.5 | 3.5 | 2.4 | 3.1 | 2.8 |
| BAC-C12 | 4.9 | 4.2 | 1.5 | 2.0 | 3.4 |
| BAC-C14 | 4.0 | 3.2 | 2.8 | 2.6 | 3.0 |
| BAC-C16 | 4.7 | 2.3 | 2.4 | 3.4 | 5.5 |
| BAC-C18 | 3.8 | 2.4 | 2.2 | 2.1 | 2.1 |
| DADMAC-C8 | 3.9 | 3.9 | 1.8 | 6.2 | 3.8 |
| DADMAC-C10 | 4.1 | 4.8 | 3.0 | 1.4 | 3.2 |
| DADMAC-C12 | 4.9 | 3.9 | 3.2 | 2.2 | 2.9 |
| DADMAC-C14 | 6.2 | 3.3 | 1.4 | 5.0 | 2.1 |
| DADMAC-C16 | 4.7 | 3.7 | 1.4 | 3.1 | 5.2 |
| DADMAC-C18 | 9.3 | 7.3 | 3.3 | 1.2 | 6.3 |
| BEC | 3.2 | 4.5 | 2.8 | 3.1 | 3.6 |
| CLQ | 3.5 | 1.4 | 3.4 | 7.5 | 6.6 |

Table S11 Retention times and MRM parameters for QACs and internal standards. Parameters for qualifier fragments are given in brackets.

| Analyte | Internal standard | RT (min) | Precursor (m/z) | Product (m/z) | DP (V) | CE (V) | Cxp (V) |
|---|---|---|---|---|---|---|---|
| ATMAC-C8 | BAC-C8-$D_7$ | 4.63 | 172 | 60 (57) | 26 (26) | 43 (31) | 6 (6) |
| ATMAC-C10 | BAC-C8-$D_7$ | 4.95 | 200 | 60 (57) | 21 (21) | 47 (35) | 6 (8) |
| ATMAC-C12 | DADMAC-C8-$D_6$ | 5.27 | 228 | 60 (57) | 21 (21) | 39 (39) | 6 (8) |
| ATMAC-C14 | DADMAC-C8-$D_6$ | 5.63 | 256 | 60 (57) | 86 (86) | 57 (43) | 8 (8) |
| ATMAC-C16 | DADMAC-C10-$D_6$ | 6.06 | 284 | 60 (57) | 91 (71) | 49 (49) | 8 (8) |
| BAC-C8 | BAC-C8-$D_7$ | 5.06 | 248 | 91 (58) | 46 (46) | 55 (37) | 8 (8) |
| BAC-C10 | BAC-C12-$D_7$ | 5.39 | 276 | 91 (58) | 11 (11) | 57 (49) | 8 (16) |
| BAC-C12 | BAC-C12-$D_7$ | 5.74 | 304 | 91 (58) | 81 (76) | 65 (65) | 8 (8) |
| BAC-C14 | BAC-C14-$D_7$ [1)] | 6.17 | 332 | 91 (58) | 76 (96) | 57 (63) | 8 (16) |
| BAC-C16 | BAC-C18-$D_7$ | 6.72 | 361 | 91 (58) | 16 (16) | 75 (63) | 8 (16) |
| BAC-C18 | BAC-C18-$D_7$ | 7.41 | 389 | 91 (58) | 26 (41) | 69 (67) | 8 (8) |
| DADMAC-C8 | DADMAC-C8-$D_6$ | 5.62 | 270 | 158 (57) | 26 (26) | 37 (47) | 10 (8) |
| DADMAC-C10 | DADMAC-C10-$D_6$ | 6.45 | 326 | 186 (57) | 141 (141) | 39 (57) | 8 (16) |
| DADMAC-C12 | DADMAC-C12-$D_6$ | 7.69 | 383 | 214 (57) | 141 (141) | 45 (63) | 8 (16) |
| DADMAC-C14 | DADMAC-C14-$D_6$ [2)] | 9.46 | 439 | 242 (57) | 146 (151) | 49 (71) | 10 (8) |
| DADMAC-C16 | DADMAC-C14-$D_6$ [2)] | 11.78 | 495 | 270 (57) | 141 (156) | 55 (81) | 10 (8) |
| DADMAC-C18 | DADMAC-C14-$D_6$ [2)] | 15.13 | 550.6 | 298 (57) | 6 (6) | 59 (99) | 14 (30) |
| BEC | BAC-C14-$D_7$ [1)] | 5.85 | 412 | 91 (320) | 61 (1) | 47 (29) | 16 (6) |
| CLQ | BAC-C8-$D_7$ | 1.38 | 122 | 58 (63) | 1 (16) | 85 (39) | 8 (12) |
| | BAC-C8-$D_7$ | 5.05 | 255.5 | 98 (58) | 131 (136) | 57 (43) | 8 (8) |
| | BAC-C12-$D_7$ | 5.73 | 311 | 98 (58) | 81 (96) | 61 (89) | 6 (18) |
| | BAC-C14-$D_7$ | 6.15 | 340 | 98 (58) | 93 (71) | 69 (60) | 8 (8) |
| | BAC-C18-$D_7$ | 7.38 | 396.1 | 98 (58) | 111 (96) | 77 (69) | 8 (6) |
| | DADMAC-C8-$D_6$ | 5.61 | 276.3 | 164 (63) | 171 (171) | 37 (53) | 8 (6) |
| | DADMAC-C10-$D_6$ | 6.44 | 332 | 192 (63) | 101 (101) | 41 (67) | 14 (14) |
| | DADMAC-C12-$D_6$ | 7.67 | 389.4 | 220 (63) | 96 (106) | 45 (45) | 8 (8) |
| | DADMAC-C14-$D_6$ | 9.44 | 444.5 | 248 (63) | 176 (176) | 51 (83) | 12 (8) |

[1)] BAC-C12-$D_7$ in case of wastewater samples
[2)] DADMAC-C12-$D_6$ in case of wastewater samples

Table S12 HPLC and MS settings

| | |
|---|---|
| HPLC | Sciex Exion LC |
| MS | Qtrap 4500 |
| Flow rate | 0.25 mL $min^{-1}$ |
| Injection volume | 20 µL |
| Mobile phase | ultrapure water + 50 mM formic acid + 10 mM ammonium formate (A), acetonitrile (B) |
| Gradient | 0 – 3 min: 0% – 75% B<br>3 – 14.5 min: 75% – 100% B<br>14.5 – 21 min: isocratic at 100% B<br>21 – 26 min: 100% – 0% B |
| Autosampler T | 15 °C |
| Column oven T | 40 °C |
| Source T | 650 °C |
| Curtain gas (CUR) | 35 |
| Collision gas (CAD) | medium |
| Ionspray voltage (IS) | 3,500 |
| Entrance potential (EP) | 10 |
| GS1 | 60 |
| GS2 | 60 |
| Dwell time per transition | 3-250 ms |

Table S13 Primer systems used for quantitative PCRs (qPCRs) and generation of qPCR standards

| Primer | 5′ - to - 3′sequence | Reference |
|---|---|---|
| 331-F | TCCTACGGGAGGCAGCAGT | Nadkarni et al. [4] |
| 518-R | ATTACCGCGGCTGCTGG | Muyzer et al. [5] |
| qacEallF | CGCATTTTATTTTCTTTCTCTGGTT | Jechalke et al. [6] |
| qacEallR | CCCGACCAGACTGCATAAGC | Jechalke et al. [6] |
| qacEallP | FAM-TGAAATCCATCCCTGTCGGTGT-TAMRA | Jechalke et al. [6] |
| intILC5_fw | GATCGGTCGAATGCGTGT | Barraud et al. [7] |
| intILC1_rv | GCCTTGATGTTACCCGAGAG | Barraud et al. [7] |
| sul1-FW | CGCACCGGAAACATCGCTGCAC | Pei et al.[8] |
| sul1-RV | TGAAGTTCCGCCGCAAGGCTCG | Pei et al. [8] |
| sul2F | TCGTCAACATAACCTCGGACAG | Byrne-Bailey et al. [9] |
| sul2R | GTTGCGTTTGATACCGGCAC | Byrne-Bailey et al. [9] |
| tetM-FW | ACAGAAAGCTTATTATATAAC | Kobayashi et al. [10] |
| tetM-RV | TGGCGTGTCTATGATGTTCAC | Kobayashi et al. [10] |
| qnrSrtF11 | GACGTGCTAACTTGCGTGAT | Marti and Balcazar [11] |
| qnrSrtR11 | TGGCATTGTTGGAAACTTG | Marti and Balcazar [11] |
| EUB9F | GAGTTTGATCMTGGCTCAG | Lane [12] |
| EUB1492R | ACGGYTACCTTGTTACGACTT | Lane [12] |
| qacEΔ1F | ATCGCAATAGTTGGCGAAGT | Sandvang et al. [13] |
| sul1B | GCAAGGCGGAAACCCGCGCC | Sandvang et al. [13] |
| pEX-for | GGAGCAGACAAGCCCGTCAGG | Eurofins/Snapgene (online) |
| pEX-rev | CAGGCTTTACACTTTATGCTTCCGGC | Eurofins/Snapgene (online) |
| pJET1.2F | CGACTCACTATAGGGAGAGCGGC | Thermo Scientific |
| pJET1.2 rv | AAGAACATCGATTTTCCATGGCAG | Thermo Scientific |

Table S14 Standards and qPCR conditions

| | 16S rRNA gene | *qacE/ qacEΔ1* | *intI1* | *sul1* | *sul2* | *tetM* | *qnrS* |
|---|---|---|---|---|---|---|---|
| Primer system (fragment length in bp) | 331-F/ 518-R | qacEallF/ qacEallR<br>qacEallP | intILC5_fw/ intILC1_rv | sul1-FW/ sul1-RV | sul2F/ sul2R | tetM-FW/ tetM-RV | qnrSrtF11/ qnrSrtR11 |
| Expected fragment size (bps) | 147 bp | 68 bp | 196 bp | 162 bp | 478 bp | 170 bp | 116 bp |
| Template DNA | *E. coli* TOP10 with pEX-A(pNORM)[#] | *E. coli* ESBL37B15-13-1E | *E. coli* TOP10 with pEX-A(pNORM)[#] | *E. coli* TOP10 with pEX-A(pNORM)[#] | *E. coli* ESBL37B15-13-1E | Manure DNA | E. coli TOP10 with pEX-A(pNORM)[#] |
| Standard fragment amplification without cloning – used primer systems | - | qacEΔ1F-sul1bR | pEX-for/pEX-rev | pEX-for/pEX-rev | - | - | pEX-for/pEX-rev |
| PCR products used for standard generation by cloning | - | - | - | - | sul2F/ sul2R | tetM-FW/ tetM-RV | - |
| Cloning vector; reamplification primer system | | - | - | | pJet1.2; pJET1.2 fw/ pJET1.2rv | pJet1.2, pJET1.2 fw/ pJET1.2rv | |
| Final standard length | 1329 | 798 bp | 1329 | 1329 | 597 | 289 | 1329 |
| Primer conc. (µM) | 0.2 | 0.3 | 0.2 | 0.2 | 0.2 | 0.2 | 0.2 |
| Probe | - | 0.25 | - | - | - | - | - |
| Standard range | $10^8$ - $10^6$,$10^4$ | $10^8$ - $10^2$ | $10^7$ - $10^1$ | $10^7$ - $10^2$ | $10^7$ - $10^1$ | $10^8$-$10^3$ | $10^4$-$10^1$ |
| Efficiency | 73 % | 94.4 % | 100.1 % | 89.4 % | 96.3 % | 96.3 % | 100.9 % |
| R^2 | 0.977 | 0.995 | 0.999 | 0.999 | 0.999 | 0.999 | 0.997 |
| Slope | -4.202 | -3.465 | -3.320 | -3.605 | -3.413 | -3.414 | -3.300 |
| y-int | 47.628 | 34.517 | 43.504 | 36.919 | 34.507 | 37.735 | 33.719 |

| | | | | | | | |
|---|---|---|---|---|---|---|---|
| qPCR program | 98°C, 2 min<br>98°C, 5 sec<br>60°C, 5 sec *<br>45 cycles | 50°C, 2 min<br>95°C, 10 min<br>95°C, 15 sec<br>60°C, 1 min*<br>40 cycles | 98°C, 2 min<br>98°C, 5 sec<br>60°C, 5 sec*<br>45 cycles | 98°C, 2 min<br>98°C, 5 sec<br>60°C, 5 sec*<br>45 cycles | 98°C, 2 min<br>98°C, 5 sec<br>60°C, 5 sec*<br>79.9°C, 5 sec*<br>45 cycles | 98°C, 2 min<br>98°C, 5 sec<br>51°C, 5 sec*<br>74°C, 5 sec*<br>45 cycles | 98°C, 2 min<br>98°C, 5 sec<br>60°C, 5 sec*<br>45 cycles |
| Melt curve | 65-95°C, 0.5°C/5 sec* | - | 65-95°C, 0.5°C/5 sec* | 65-95°C, 0.5°C/5 sec* | 65-95°C, 0.5°C/5 sec* | 65-95°C, 0.5°C/5 sec* | 65-95°C, 0.5°C/5 sec* |

# [14]

* Reading

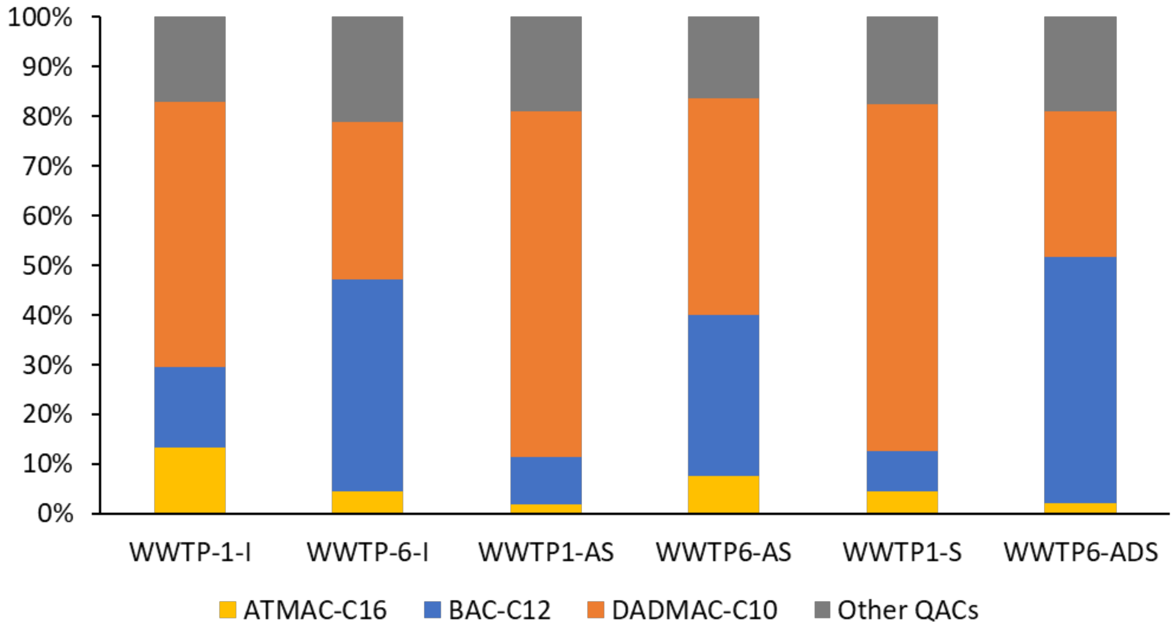


Fig. S1 Proportion of ATMAC-C16, BAC-C12, and DADMAC-C10 in the total QAC concentrations (excluding DADMAC-C18 and CLQ)

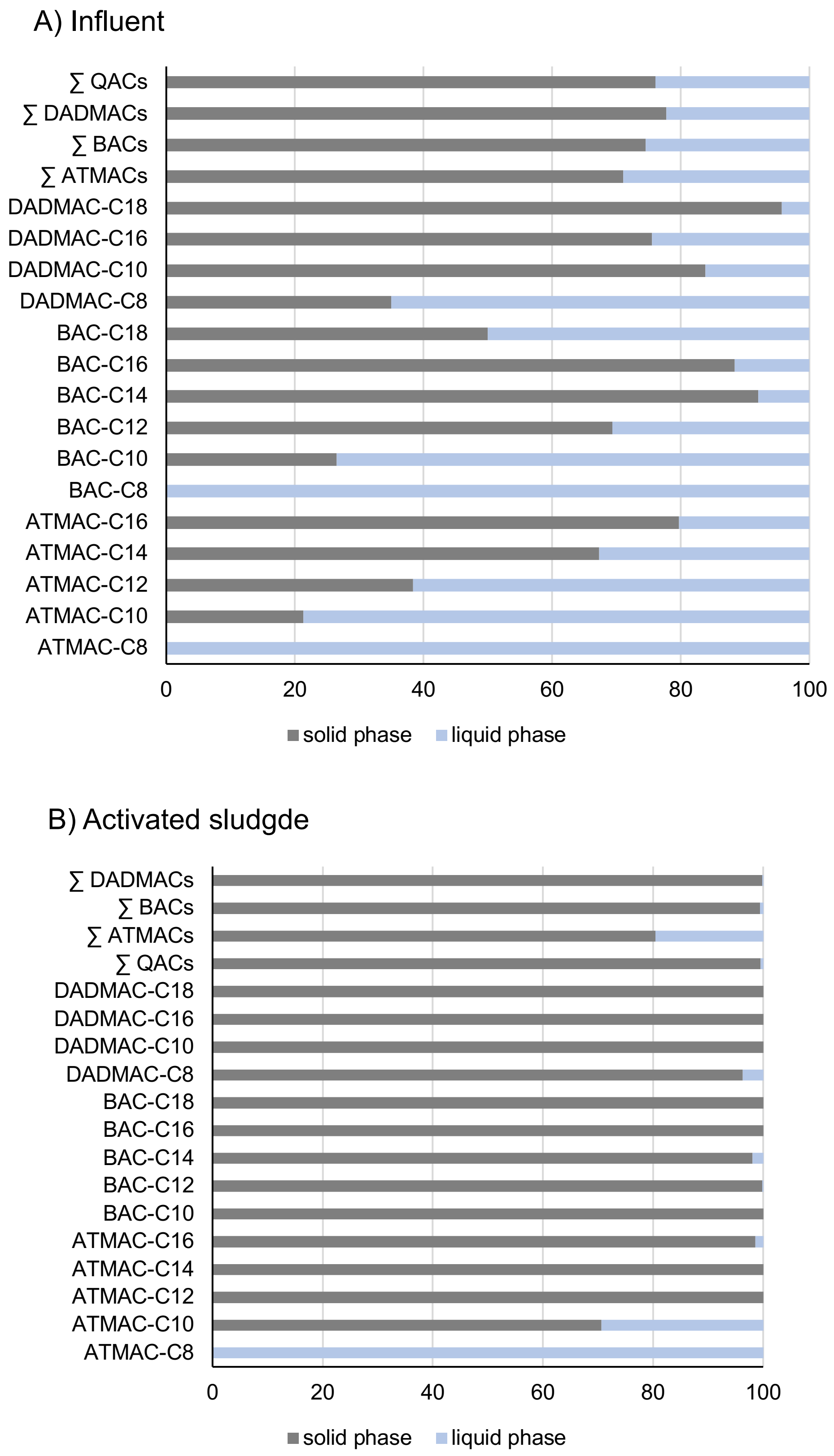


Fig. S2 Phase distribution of QACs in influent (A, n=3) and activated sludge (B, n=4). Values were averaged over all investigated samples and are only shown for homologues detected in at least one phase. Concentrations <LOD were treated as 0.

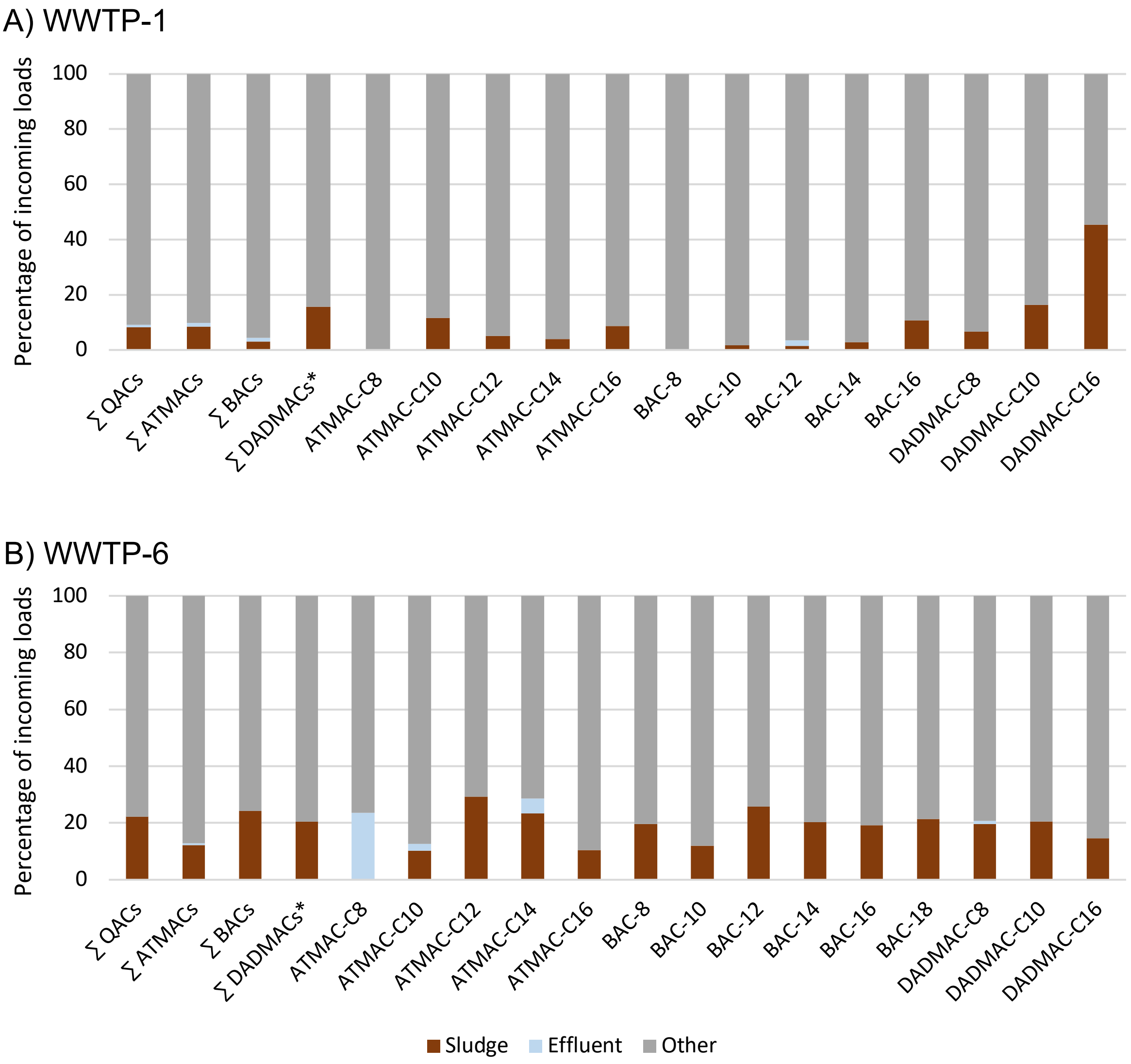


Fig. S3 QAC mass balances in the rural (A) and urban WWTP (B). The fractions were estimated based on the measured QAC concentrations and annual sludge production and processed wastewater volumes. “Other” is the unaccounted fraction which includes transformation and removal via biodegradation.

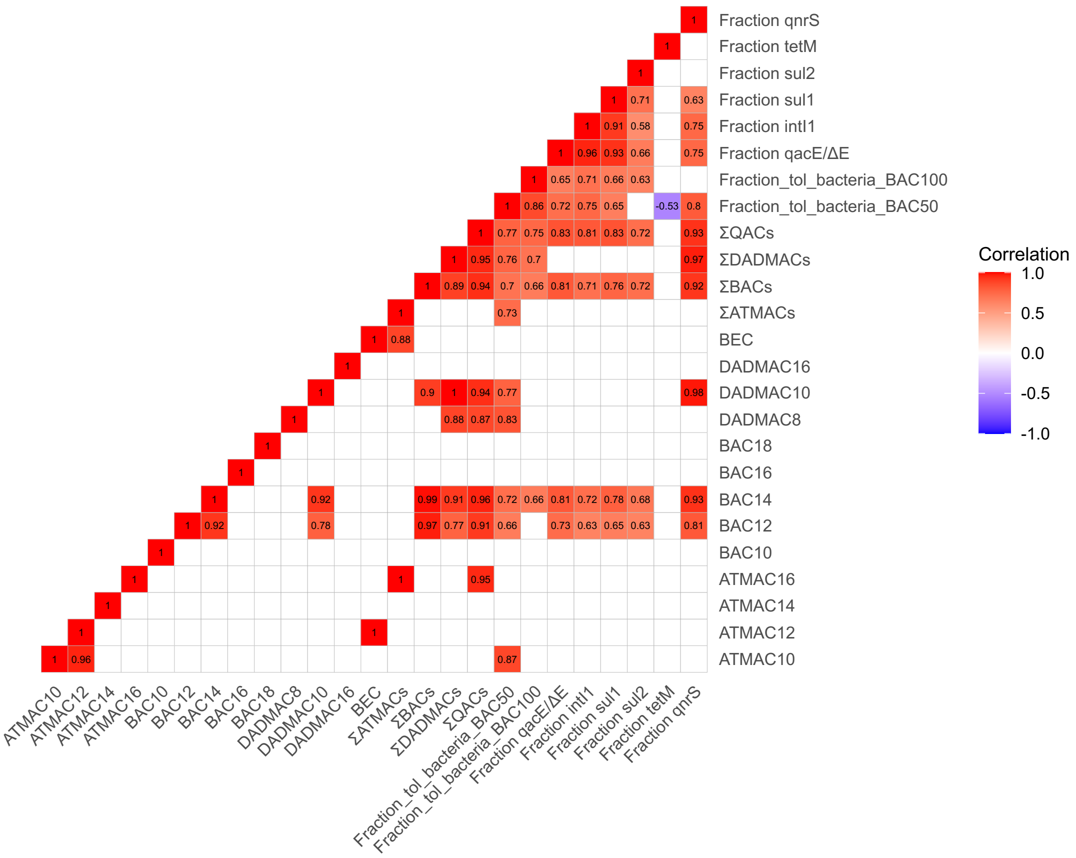


Fig. S4 Spearman correlations between dry weight concentrations of QACs, the fraction of BAC-C12 tolerant bacteria, and relative gene abundances among all analysed (semi-)solid samples (manure, biogas plant digestate, activated sludge, dewatered sludge and anaerobically digested sludge; n=30). Only significant correlation coefficients are shown ($p_{adjusted} > 0.05$).

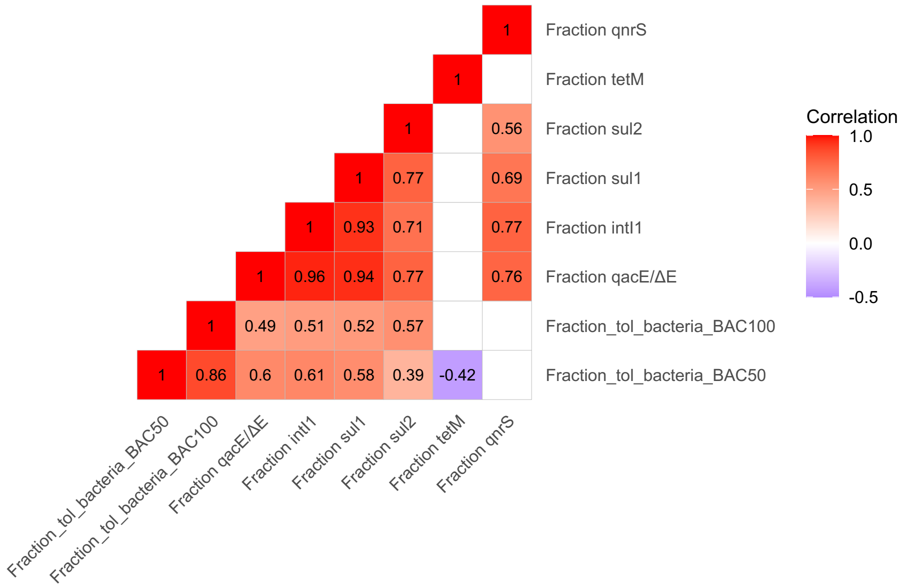


Fig. S5 Spearman correlations between the fraction of BAC-C12 tolerant bacteria and relative gene abundances among all analysed samples (manure, biogas plant digestate, activated sludge, dewatered sludge, anaerobically digested sludge, WWTP influent, WWTP effluent; n=37). Only significant correlation coefficients are shown ($p_{adjusted} > 0.05$).

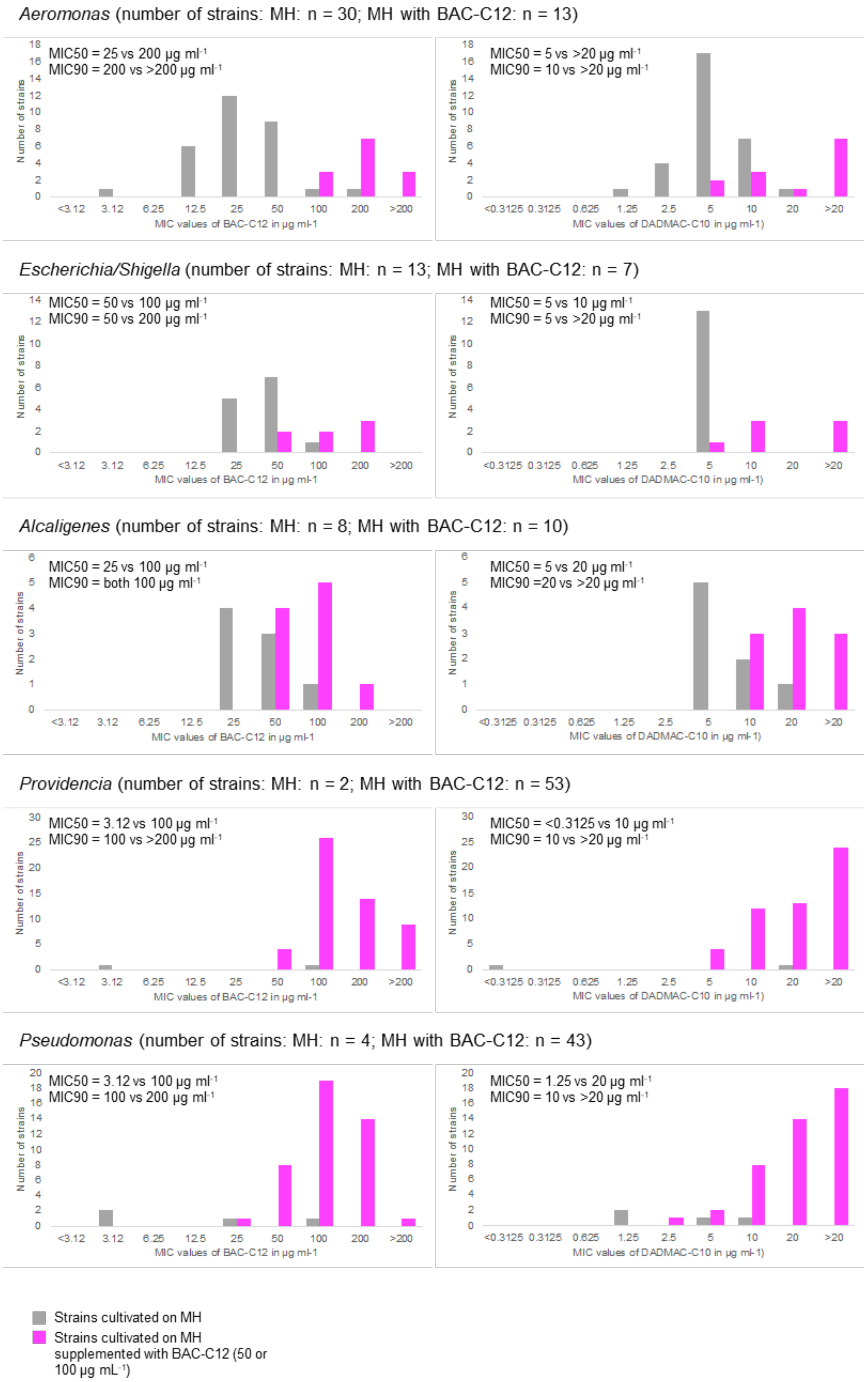


Fig. S6 Comparative analysis of BAC-C12 and DADMAC-C10 MIC distribution patterns among strains of the three most abundant genera cultivated in the absence and presence of BAC-C12 (*Aeromonas, Escherichia/Shigella,* and *Alcaligenes*). MIC50 and MIC90 values are given for strains of individual genera.

*Providencia* (number of strains: M: 10; BGP-D: 10; AS: 3; S: 11; ADS: 1; I: 12; E: 6)

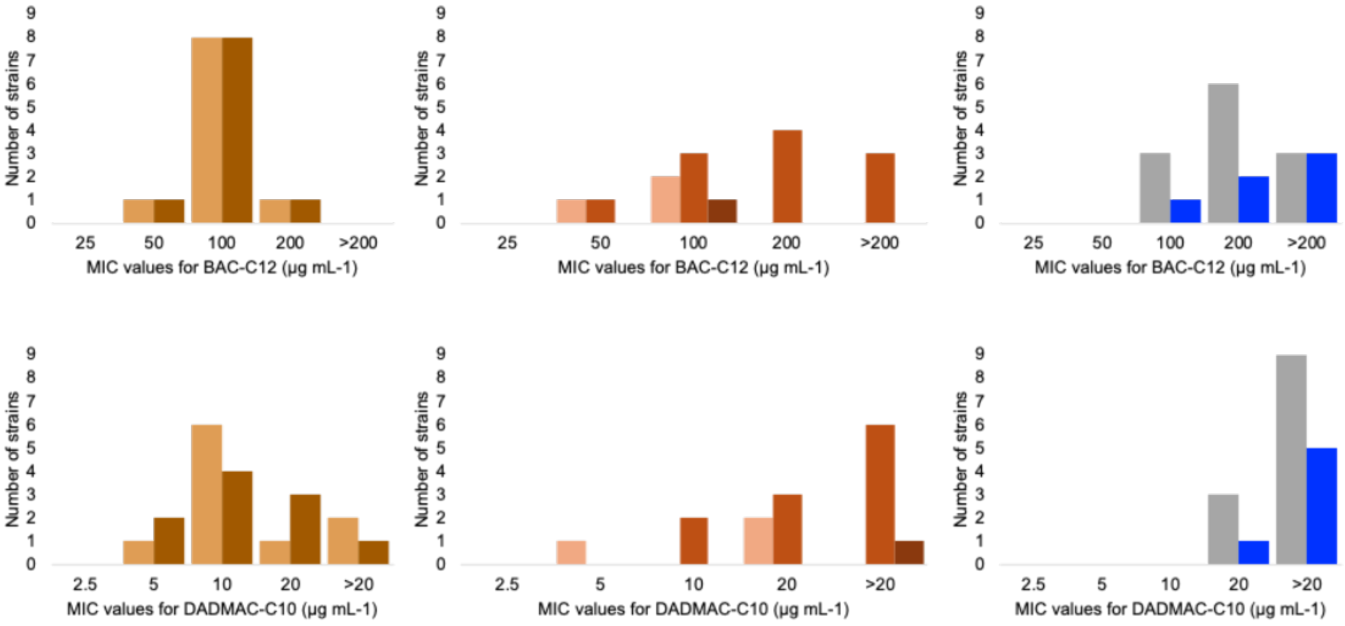


*Pseudomonas* (number of strains: M: 21; BGP-D: 5; AS: 4; S: 4; ADS: 5; I: 4; E: 4)

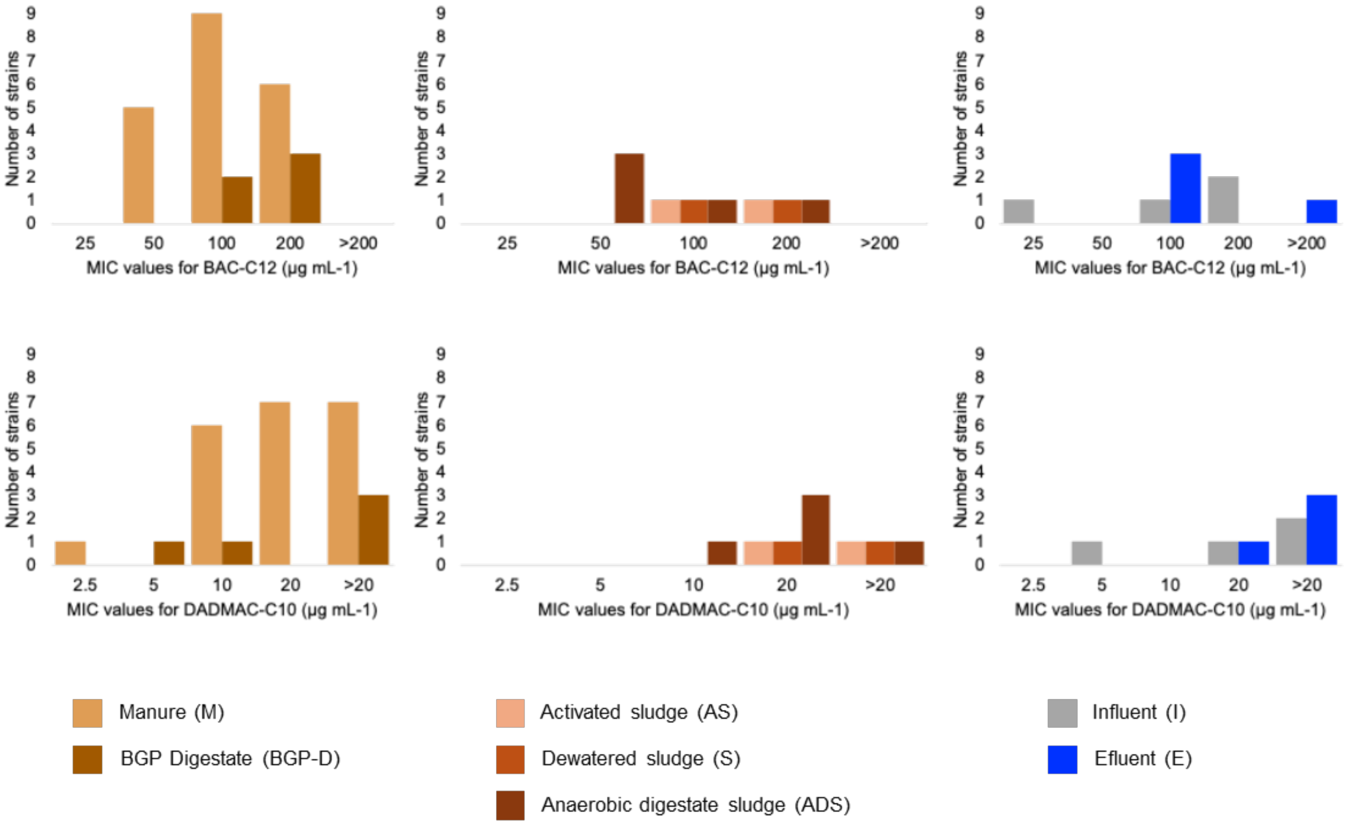


Fig. S7 Comparative analysis of BAC-C12 and DADMAC-C10 MIC distribution patterns for *Providencia* and *Pseudomonas* strains separated based on individual sampling types of both waste stream systems.
